# Harbour seals are regaining top-down control in a coastal ecosystem

**DOI:** 10.1101/267567

**Authors:** Geert Aarts, Sophie Brasseur, Jan Jaap Poos, Jessica Schop, Roger Kirkwood, Tobias van Kooten, Evert Mul, Peter Reijnders, Adriaan D. Rijnsdorp, Ingrid Tulp

## Abstract

Historic hunting has led to severe reductions of many marine mammal species across the globe. After hunting ceased, some populations have recovered to pre-exploitation levels, and may again act as a top-down regulatory force on marine ecosystems. Also the harbour seal population in the international Wadden Sea grew at an exponential rate following a ban on seal hunting in 1960’s, and the current number ∼38,000 is close to the historic population size. Here we estimate the impact of the harbour seal predation on the fish community in the Wadden Sea and nearby coastal waters.

Fish remains in faecal samples and published estimates on the seal’s daily energy requirement were used to estimate prey selection and the magnitude of seal consumption. Estimates on prey abundance were derived from demersal fish surveys, and fish growth was estimated using a Dynamic Energy Budget model. GPS tracking provided information on where seals most likely caught their prey.

Harbour seals from the Dutch Wadden Sea fed predominantly on demersal fish, e.g. flatfish species (flounder, sole, plaice, dab), but also sandeel, cod and whiting. Total fish biomass in the Wadden Sea was insufficient to sustain the estimated prey consumption of the entire seal population year-round. This probably explains why seals also acquire prey further offshore in the adjacent North Sea, only spending 13% of their diving time in the Wadden Sea. Still, seal predation was estimated to cause an average annual mortality of 43% and 60% on fish in the Wadden Sea and adjacent coastal zone, respectively. There were however large sources of uncertainty in the estimate, including the migration of fish between the North Sea and Wadden Sea, and catchability estimates of the fish survey sampling gear, particularly for sandeel and other pelagic fish species.

Our estimate suggested a considerable top-down control by harbour seals on demersal fish. However predation by seals may also alleviate density-dependent competition between the remaining fish, increasing fish growth, and partly compensating for the reduction in fish numbers. This study shows that recovering coastal marine mammal populations could potentially become an important component in the functioning of shallow coastal systems.

## INTRODUCTION

Large-scale historic whaling and sealing led to a severe global decline of many marine mammal species (Clapham et al. 1999, Baker and Clapham 2004). As a consequence, marine ecosystems may have lost important regulating forces from such top-predators (Heithaus et al. 2008, Estes et al. 2016). While some marine mammal populations have not fully recovered after hunting ceased (Baylis et al. 2015), or have even continued to decline (Springer et al. 2003), others have gone through rapid increases (Brasseur et al. 2015, 2018) reaching or exceeding presumed pre-exploitation levels (Roman et al. 2015). This raises the question how such recoveries influence the food web regulation in marine ecosystems.

Harbour seals (*Phoca vitulina*) have been top predators in the Wadden Sea of the Netherlands, Germany and Denmark, since its formation 7,500 year ago. Due to severe human hunting, their numbers declined from an estimated 40,000 in 1900 to approximately 4,500 around 1960 (Reijnders 1992). After a ban on seal hunting, the population grew at an annual rate of 12% (Brasseur et al. 2018). Currently, the population is approaching pre-1900 levels with approximately 38,000 individuals regularly hauling out in the international Wadden Sea (Galatius et al. 2017), of which approximately 10,000 in the Dutch part of the Wadden Sea (Figure 1).

**Figure 1.**
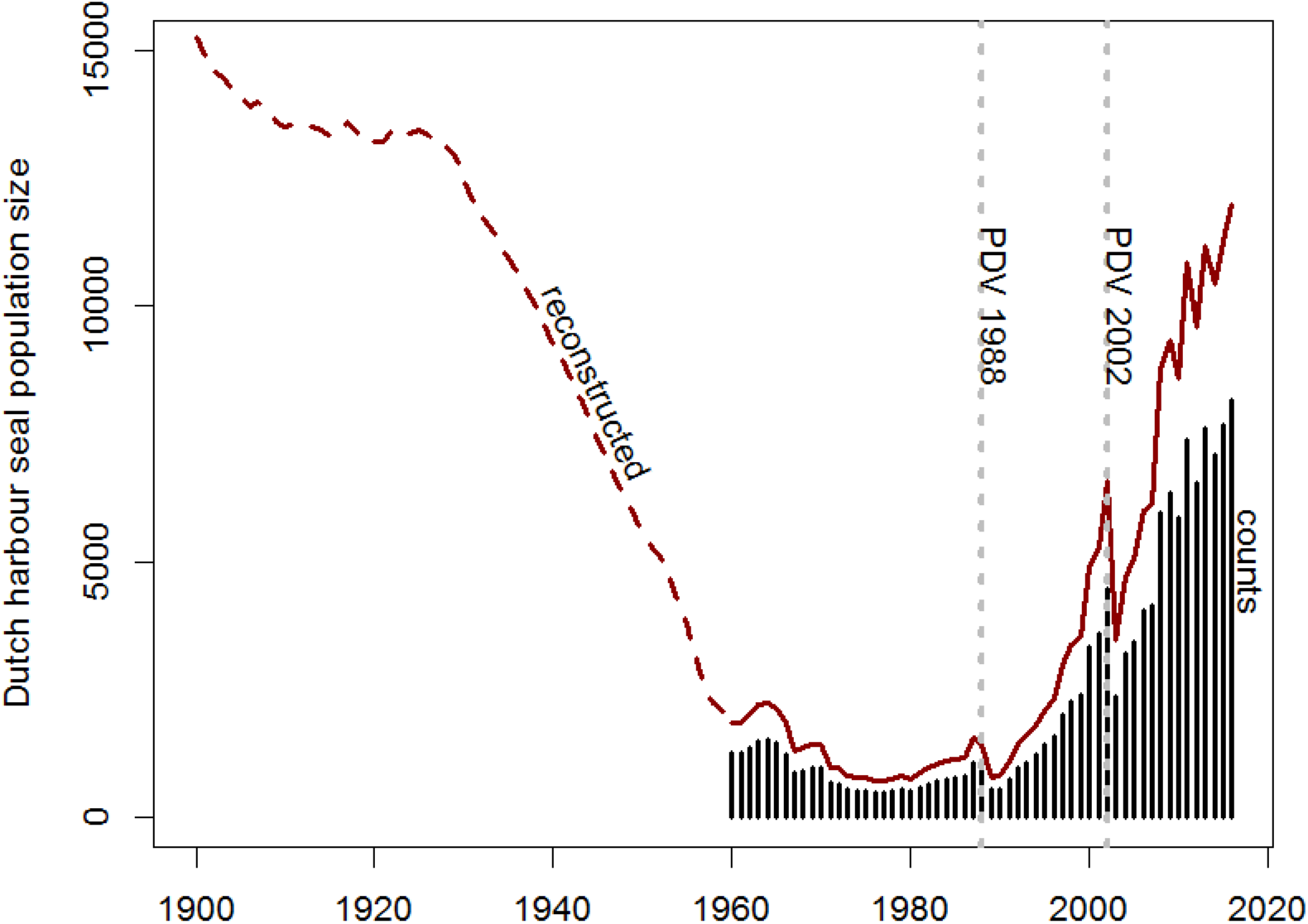
Development of the harbour seal population in the Dutch Wadden Sea. The maximum numbers of seals observed during the moult survey are represented by the vertical bars(Brasseur et al. 2018). Prior to 1960, the estimated population size was reconstructed based on seal hunting statistics (Reijnders 1992), while from 1960 onwards, the population size was estimated to be 1.47 (i.e. 0.68^−1^) times the observed counts, to correct for animals at sea at the time of the census (Ries et al. 1998). In 1988 and 2002, the population was reduced by approximately 50% due to the Phocine distemper virus (PDV).

Harbour seals require approximately 4-5 kg of fish each day, although this varies by season and the seals’ size (Härkönen and Heide-Jørgensen 1991). They are considered generalist predators feeding on both demersal and pelagic fish species, but in shallow, soft-sediment regions like the southern North Sea, they mostly eat flatfish (∼75%) such as flounder (*Platichtys flesus*), sole (*Solea solea*), and dab (*Limanda limanda*), but also gadoids, sandeel (*Ammodytidae*) (Härkönen 1987, Tollit et al. 1997, Kavanagh et al. 2010).

The strong growth of the harbour seal population in recent decades occurred almost at the same time as the abundances of their prey in the Wadden Sea and the adjacent coastal zone of the North Sea declined (Tulp et al. 2008, 2017). Some size classes, such as >1 year old plaice and flounder, have almost completely disappeared from the Wadden Sea (van Keeken et al. 2007, van der Veer et al. 2011). Several hypotheses have been put forward to explain the declines in the fish biomass, e.g. increasing water temperature, increased human activities, declining nutrients due to cleaner river outflows, fishery bycatch or increased predation by birds (e.g. cormorants) and seals (Temming and Hufnagl 2015, van der Veer et al. 2016, Tulp et al. 2017).

The objective of this study is to estimate the role of predation pressure by harbour seals on the demersal fish community in the Dutch Wadden Sea and nearby coastal North Sea. The specific value of this study is that such estimates of predation pressure can subsequently be included as an additional source of mortality into fish population models (Tyrrell et al. 2011). More generally, this study may shed light on how marine coastal ecosystems function under the presence of large numbers of marine top-predators.

## METHODS

### Analysis structure

We estimated the impact of seals on the local fish abundance as follows:

1. Harbour seal diet was defined based on analysis of faecal samples collected on haul-out sites in the Dutch Wadden Sea.
2. Based on estimated seal energy requirements, and energetic content of the prey in their diet, the average daily fish consumption was estimated.
3. The total daily fish consumption by all harbour seals was estimated, taking into account the size of the population in the Dutch Wadden Sea and the time spent foraging in different regions (using data from GPS tracked harbour seals).
4. Total number and biomass of prey species present in three regions (Wadden Sea, Wadden coastal zone up to 25 m depth (‘Wadden coast’) and offshore North Sea up to 50 km from the nearest haul-out) were estimated using data from the annual demersal fish survey (DFS) and beam trawl survey (BTS) collected in September
5. Growth in prey biomass from September onwards was reconstructed using a dynamic energy budget (DEB) model. The model was fitted to seasonal length distribution data, and accounted for daily variations in temperature.
6. The reconstructed prey numbers and biomass were subsequently reduced by the estimated daily food intake by all harbour seals.

### 1. Harbour seal diet based on faecal analysis

Between 2002 and 2009, 103 faecal samples were collected opportunistically from tidal haul-out sites throughout the Dutch Wadden Sea. Most samples were collected near Texel in the west (i.e. ‘Noorderhaaks’ and ‘Steenplaat’) and near Schiermonnikoog in the east (i.e. ‘Simonszand’) (Figure 2a). After collection, samples were frozen (−20°C) until further processing. For the analysis, samples were placed in a meshed (120μm) bag and washed in a laundry washing machine at 70°C, including a prewashing cycle using a biological detergent (Biotex, Unilever). Samples were then dried. Recognisable fish remains (otoliths and bones) were picked out using a binocular microscope. These hard parts were identified using available reference guides (Härkönen 1986, Watt et al. 1997) and compared to a reference collection. For this study only otoliths were used. Otoliths were measured using digital callipers (to 0.1 mm), and assigned to a wear class. After correction for wear (Leopold 2015), both otolith length and width were used to infer fish length based on known otolith length - fish length regressions (Härkönen 1986, Watt et al. 1997, Leopold et al. 2001). Fish weights were inferred using known fish length – weight regressions ((Robinson et al. 2010) and WMR unpublished data). The ten most important species in the diet were identified based on calculated contributions by fresh mass to the seal diet.

**Figure 2a. Top figure:**
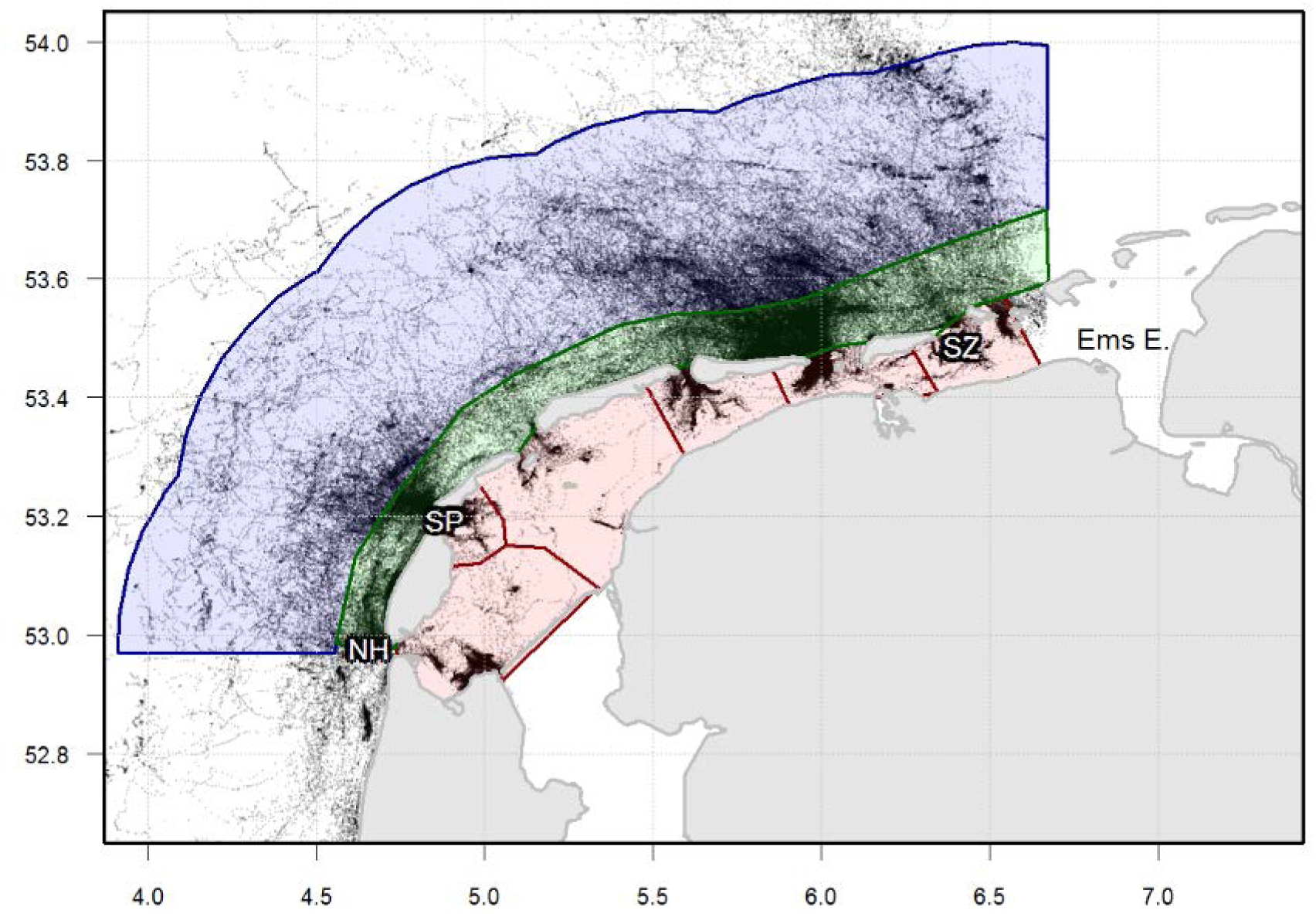
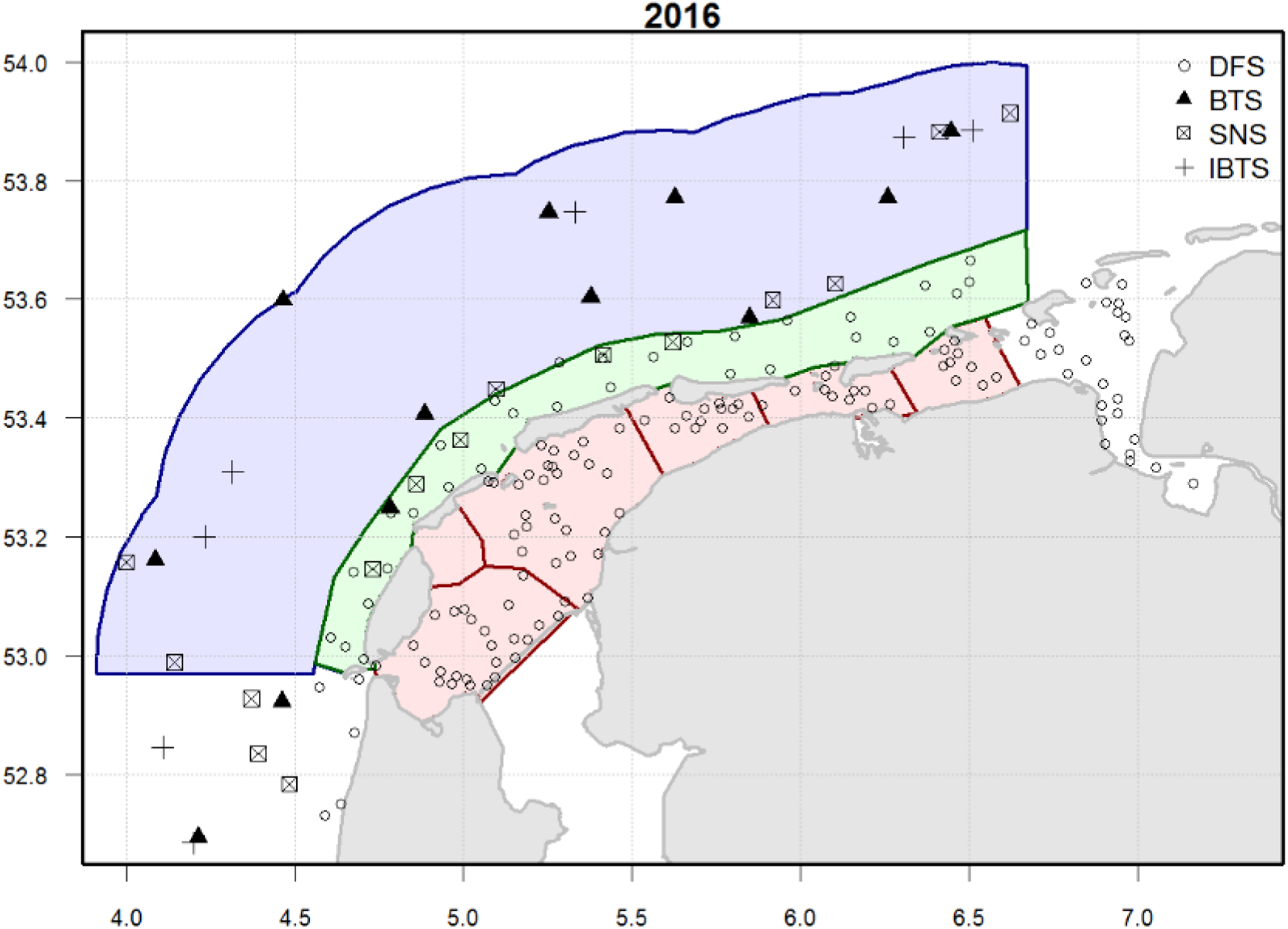
Distribution of harbour seals, and abbreviations of some faecal sampling locations. Data from the 149 tracked harbour seals making trips from haul-out sites located in the Wadden Sea. Trips of seals making trips from haul-out sites located the Ems Estuary (Ems. E), were excluded. Most faecal samples were collected on the ‘Noorderhaaks’ (NH), ‘Steenplaat’ (SP) and ‘Simonszand’ (SZ). **b. Bottom figure**: Fish survey areas along the North Sea coasts of The Netherlands (adapted from Tulp et al. 2016). Harbour seal fish consumption was assessed in three zones: the **Wadden Sea** (shaded pink – corresponding to ICES areas 610, 612, 616-619), **Wadden coast** (shaded lime green – ICES area 404) and a **North Sea offshore zone** up to 50km away from the seal haul-out sites located in the Dutch Wadden sea (shaded blue). The points represent fish survey locations from 2016.

### 2. Harbour seal average daily food requirement

Harbour seals require energy for maintenance and growth (Markussen et al. 1990), reproduction (i.e. foetal growth, lactation, mating), moult, and activities such as locomotion. Juvenile phocid seals require on average 1.4 times more energy per kg body weight for maintenance and growth than adults (Innes et al. 1987), but have a lower overall energy requirement because they are smaller in size and do not take part in reproduction. A study on captive harbour seals (four adult males, one sub adult male and 1 adult female) which were fed ad libitum, the gross energy intake was estimated to be 6071 kcal/day (Rosen and Renouf 1998). These estimates were close to the estimates for adult males used by (Härkönen and Heide-Jørgensen 1991, i.e. 5890 kcal/day). When taking the local population structure into account, Härkönen & Heide-Jørgensen (1991) estimated that on average harbour seals in the Skagerrak (Sweden) have an ingested energy requirement (*ER*) of 4680 kcal per day.

To meet this energy requirement, Härkönen & Heide-Jørgensen (1991) estimated that harbour seals should consume 4.1 kg of flatfish each day in the Skagerrak. The energy densities of their prey, and hence also the seals’ food intake varies between prey species and seasons (Pedersen and Hislop 2001). The energy density of fish is often highest just after the growth season, prior to the winter, which is subsequently devoted to maintenance and reproduction (Dawson and Grimm 1980). Whenever possible, we used caloric densities from late summer, early winter (see Table 1). The average daily consumption *C* (in kg) per seal, was estimated based on the energy required (ER) and species-specific caloric densities (*ED* in kcal/g), weighted by the relative occurrence of prey *W*_*i*_ in the harbour seal diet:

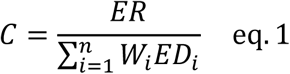

### 3. Total fish consumption and distribution of foraging effort

To estimate the total fish consumption by harbour seals in the Wadden Sea and nearby waters, information is needed on how many seals there are, and where they acquire their food. Harbour seal count data were collected annually in the Dutch Wadden Sea using aerial surveys since the 1960s (Brasseur et al. 2018). To correct for the number of seals in the water during the aerial survey counts, this studied assumed a haul-out probability of 68% (Ries et al. 1998).

To study the seals’ distribution at sea, we relied on data from animal-borne GPS data loggers collected in a series of research projects. In total, 225 harbour seals were tracked in the Netherlands between 2007 and 2015, with most (142 individuals) tracked from the Ems estuary (tagged between 2009 and 2011). Harbour seals were caught on haul-out sites with a large seine net, fitted with Fastloc GPS data loggers glued to the fur of the neck using epoxy, and released directly on location. Loggers fell off as hair weakened during the annual moult (July-August), if they did not dislodge before-hand. The GPS data loggers also contained depth and submergence sensors: These were used to determine the activity of the seal: “diving” (deeper than 1.5 m for at least 8 s), “at surface” (no dives for 180 s) or “hauled out” (start = continuously dry for at least 600, end = wet for at least 40 s). Dive records included a description of the dive-profile at 23 points of the dive, but also summary data on maximum dive depth, dive duration, and surface interval duration. All data were stored on-board and transmitted, when in contact with a GSM base.

The seal location data from the GPS loggers were classified into trips, where the start and end time of each trip was defined by the haul-out data from the loggers. The seal’s haul-out location was linked to the nearest known haul-out site (based on aerial survey data). Only locations from trips starting at haul-out sites in the defined study area (i.e. the Dutch Wadden Sea, see Figure 2) were used. Location data from seals making trips from haul-out sites located within the Ems estuary were excluded, since they were over-represented in the sample, and inclusion would result in an over-estimate of the time harbour seals spent within the Wadden Sea. The seal GPS location data were used to estimate the fraction of time spent at sea, within three zones: i.e. the Wadden Sea, the adjacent coastal zone up to the 25 m depth contour (‘Wadden coast’) and the offshore area between the 25 m depth contour up to 50 km distance from haul-out sites (Figure 2). The proportion of time spent ‘foraging’ within the different regions was estimated by calculating the dive time (i.e. time spent below 1.5 m depth) in each region, as fraction of total dive time. It was not possible to identify feeding dives from other dives (e.g. resting or transiting), because Wadden Sea and nearby waters are shallow (max 30 m), and the bottom is always within reach.

### 4. Estimation of total number of biomass prey species present in Wadden Sea and nearby waters

Data were available from four fish surveys; the demersal fish survey (DFS), the Dutch beam trawl survey (BTS), the international bottom trawl survey (IBTS) and the sole net survey (SNS) (see Figure 2b for distribution of sample locations, see Appendix 2: Table S1 describing the gear characteristics). The DFS has been conducted annually in September-October since 1970. The DFS covers the Wadden Sea and coastal waters (up to 25 m depth) from the southern border of the Netherlands to Esbjerg (van Beek et al. 1989). A smaller beam size (3 m) was used in the Wadden Sea, to navigate the often-narrow gullies, while a larger, more robust beam (6 m) was used in the more exposed adjacent waters. Both gears were rigged similarly, only the size of the beam differed: the trawls were rigged with one tickler chain, a bobbin rope, and a fine-meshed cod-end (20 mm). In the Wadden Sea, fishing was restricted to the tidal channels and gullies deeper than 2 m because of the draught of the research vessel. By fishing at low speed (2–3 knots) and using a fine meshed cod end (20 mm), larger sized fish (>15 cm) were relatively underrepresented in the surveys. The gear used is suitable for demersal species, but suboptimal for pelagic species such as herring and sprat. Totals of 110-120 hauls in the Wadden Sea and 30 hauls in the adjacent coastal zone were taken annually, unless adverse weather conditions limited sampling. During each haul the position, date, time of day and depth were recorded. Fish species were identified and measured to the nearest cm. Local fish densities (n/10,000 m^2^) were calculated from fish counts per haul using the travelled distance during the haul and the beam width to calculate the swept area.

The Dutch BTS covers the central North Sea, and is designed to sample the older flatfish species (i.e. ≥1 year old). Compared to the DFS, the BTS is carried out with a larger beam trawl (8 m), a higher speed (4 knots) and a larger mesh size of 120 mm, with 40 mm stretched mesh cod end (Rogers et al. 1998). Another fish survey in the region is the IBTS (Heessen and Daan 1996). Its aim is to obtain recruitment-indices for species, like cod, haddock, whiting, Norway pout, mackerel, saithe, herring and sprat, but catchability of flatfish species is very low. The SNS focuses on 1-4 year old sole (van Keeken et al. 2007), but fishing only takes place at pre-defined transects parallel or perpendicular to the coast. Because of the above mentioned limitations, this study only uses the DFS for the Wadden Sea and nearby coastal zone, and the BTS for regions further offshore regions (up to 50 km from the nearest Wadden Sea seal haul-out site). Only data from quarter 3 were used.

In most fish surveys, a variable fraction of the fish present in the path of the net will be caught (i.e. incomplete catchability), which depends on the vertical distribution of fish in the water column and gear efficiency. Gear efficiency depends on a number of factors, such as the gear-type used (e.g. mesh size, net configuration), fishing speed, fish length, fish behaviour, water clarity and fish species (Dickson 1993). Catchability estimates of the DFS are unavailable for demersal species, but Kuipers (1975) attempted to unravel how several processes influence the catching efficiency for plaice. The data provided in (Kuipers 1975) were used to re-estimate the catching efficiency functions based on more up-to-date statistical techniques. In addition, we also accounted for escape underneath the net based on (Reiss et al. 2006). See Appendix S1 for a detailed description of the analysis and results. For the 3m beam trawl used by the DFS, we arrived at an overall catching efficiency which closely corresponds to the efficiency provided by Bergman *et al.* (1989).

To determine the catching efficiency of the BTS, SNS and IBTS, the catching efficiency for the DFS in the Wadden coast was used as a reference, and the DFS catchability was subsequently corrected by comparing length-specific numbers per unit of effort between the DFS and other surveys (see Appendix S1). Given the lack of catchability estimates for any of the other demersal fish species, we assumed them to be similar to plaice.

### 5. Fish growth based on Dynamic Energy Budget models

The seal consumption was calculated in kg/day. To convert this into the *number* of fish caught each day, it was necessary to know the average weight of fish. Since fish grow, seasonal changes in weight need to be taken into account. Growth is species specific, and depends on fish size, ambient temperature, and food conditions. These aspects can be parameterized in a Dynamic Energy Budget (DEB) model (Sousa et al. 2010). Here, DEB models were used for the five important prey species, for which DEB parameters were readily available, namely flounder, plaice, dab, sole and bull-rout. DEB model specification and parameters were based on (van der Veer et al. 2009, Freitas et al. 2010, Teal et al. 2012). The volumetric daily growth (in gram) was defined as

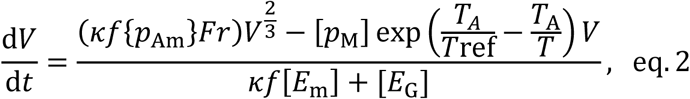

*V* is the weight of the individual fish (in gram) and *T* is ambient water temperature (in K). κ is the fraction of utilized energy spent on maintenance plus growth, *f* is a multiplication factor for food availability (*f*=1, ad libitum) and {*p*_Am_} is maximum surface area specific assimilation rate (in J cm^−2^d^−1^). [*p*_M_]is the volume-specific maintenance cost, which is based on field data and hence also includes the cost for feeding and activity (e.g. swimming) (van der Veer et al. 2009). *T*_A_ is Arrhenius temperature (in K), *T*_ref_ is the reference temperature at which the assimilation rate is known, [*E*_m_] is the maximum storage density (J cm^−3^), and [*E*_G_] and is the volume-specific costs of structure (J cm^−3^). *Fr* is the enzyme fraction that is in its active state, which is temperature dependent (see eq. 2 in (van der Veer et al. 2009)). Here for all species *Fr* was based on plaice, given the lack of the necessary parameters for the others species.

Preliminary runs of our DEB models suggested that the reduced growth in late summer (also observed in van der Veer et al. 2016) and winter could not be explained by low temperature alone (Appendix S2: Figure S2). To parameterize the DEB model in order to account for additional seasonal variability in growth, field data on seasonal variation in fish length were used. The fish length data were collected during fish surveys of the National Programme Sea and Coastal Research (van der Veer et al. 2016). Sampling took place in the western Wadden Sea in 2009 and 2010 during most months, except for the winter months (December to February), and covered both the intertidal areas (using 2-m beam trawl towed from a rubber dinghy) and subtidal areas (using a 3-m beam trawl towed by the 20m, low draft vessel *RV Navicula*). Sampling took place during day-time, and was centred around high tide (i.e. 3 hours before and after high tide). See (Freitas et al. 2016) for more details.

The DEB model was fitted to monthly mean length estimates of 0-and 1-year old fish (Figure 6, Appendix S2: Figure S3a-e). While all parameters were fixed (see, van der Veer et al. 2009, Freitas et al. 2010, Teal et al. 2012), the food availability parameter *f* was estimated, and allowed to vary between summer (*f*_*summer*_) and winter (*f*_*winter*_), and also the onset of summer (*t*_*summer*_) and winter (*t*_*winter*_) was estimated (based on least-squares in R-function ‘optim’).

The final parameterized DEB models for the five species were used to estimate the average daily growth, taking the size distribution, ambient water temperature and estimated food availability into account. The initial size distribution was based on the abundance and size distribution of each species measured during the DFS surveys in September surveys (Tulp et al. 2017). Temperature data were obtained from the website http://live.waterbase.nl, for the station “Den Helder veersteiger” (52.96328 º N, 4.77805 º E). This procedure allowed for the estimation of total biomass and the reconstruction of biomass growth.

### 6. Reducing fish numbers and biomass by harbour seal consumption

During the survey in September (*t*=0), after correcting for catchability, the DFS and BTS provide an estimate of both biomass *B* and total number of fish *N* present in each region *j*. Each day, the biomass can increase as a result of fish growth (defined by the variable *v*). The fish growth was predicted by the DEB model, which took the water temperature, seasonal variation in food density and initial length distribution of fish caught during the survey into account (see also eq. 2). The biomass can also be reduced by seal predation. Particularly after the moult, in September, harbour seals are presumed to feed intensively to replenish losses in condition experienced during the reproduction and moult-period, and build up a fat layer for the winter (Kastelein et al. 2005).

We assumed that intake rate increases linearly with food density (type I functional response), i.e. the time spent foraging in areas with higher food density leads to a higher intake, and hence a larger reduction in biomass. Therefore, the proportion of dive time *p* in each region *j*, was weighted by food density 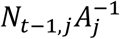 (where *A*_*j*_ is the area of the region *j*), to arrive at the proportional intake 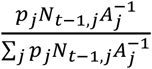. This was subsequently multiplied by the total number of seals *S*, and average daily consumption (*C*).

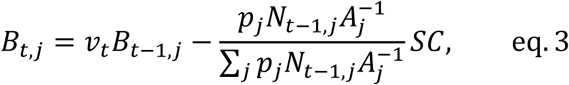

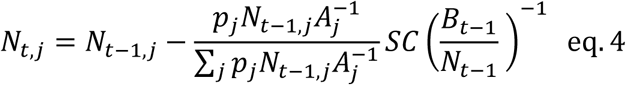

Similarly, the changes in numbers of prey fish *N* were estimated, with the main difference that fish numbers only increased due to recruitment. Here, 0-year old fish were assumed to ‘recruit’ in September, around the time of the survey. To estimate the daily food requirement in numbers, the average daily food intake (in kg) was divided by the average weight of the remaining fish (i.e. *B*_*t*-1_/*N*_*t*-1_).

The uncertainty in the estimated impact of seal predation on fish numbers and biomass is influenced by a large number of parameters, for which the uncertainties are often unknown. One of the largest uncertainties is expected to occur in the catchability; e.g. a change in catchability from 20% to 10% would lead to a doubling of the number of fish present. The uncertainty in our catchability estimate was caused by the uncertainty in the proportion of lateral escape, avoidance of the approaching vessel, escape underneath, and for the BTS, also the translation from the DFS-to BTS-catchability based on length-specific catch ratios (Appendix S1). Each step in this chain produces a (length-specific) selectivity curve quantified by parameters and corresponding uncertainties. We repeatedly sampled (500 times) from these parameter distributions, to estimates how uncertainty in the catchability propagates into the estimated effect of seal predation (eq. 3 and 4). Species-specific effects on catchability were not taken into account.

## RESULTS

### Seal diet and daily consumption

In total, 79 faecal samples contained otoliths (n= 2168). Of these, 36 samples were collected in September, 17 in August, and the rest in November (8), March (5), April (5), January (3), October (2), December (2) and July (1). Samples were collected between 1999 and 2009, with most samples from 2005 (29), 2007 (26) and 2002 (10). The 10 most represented fish species (based on estimated fresh weight) were flounder (39% of the total estimated weight), sandeel (17%), sole (14%), five-bearded rockling (5.1%), whiting (4.6%), plaice (4.4%), cod (4.3%), dragonet (2.1%), dab (2%) and bull-rout (2%) (Figure 3). Together, these prey represent 95% of the diet.

**Figure 3.**
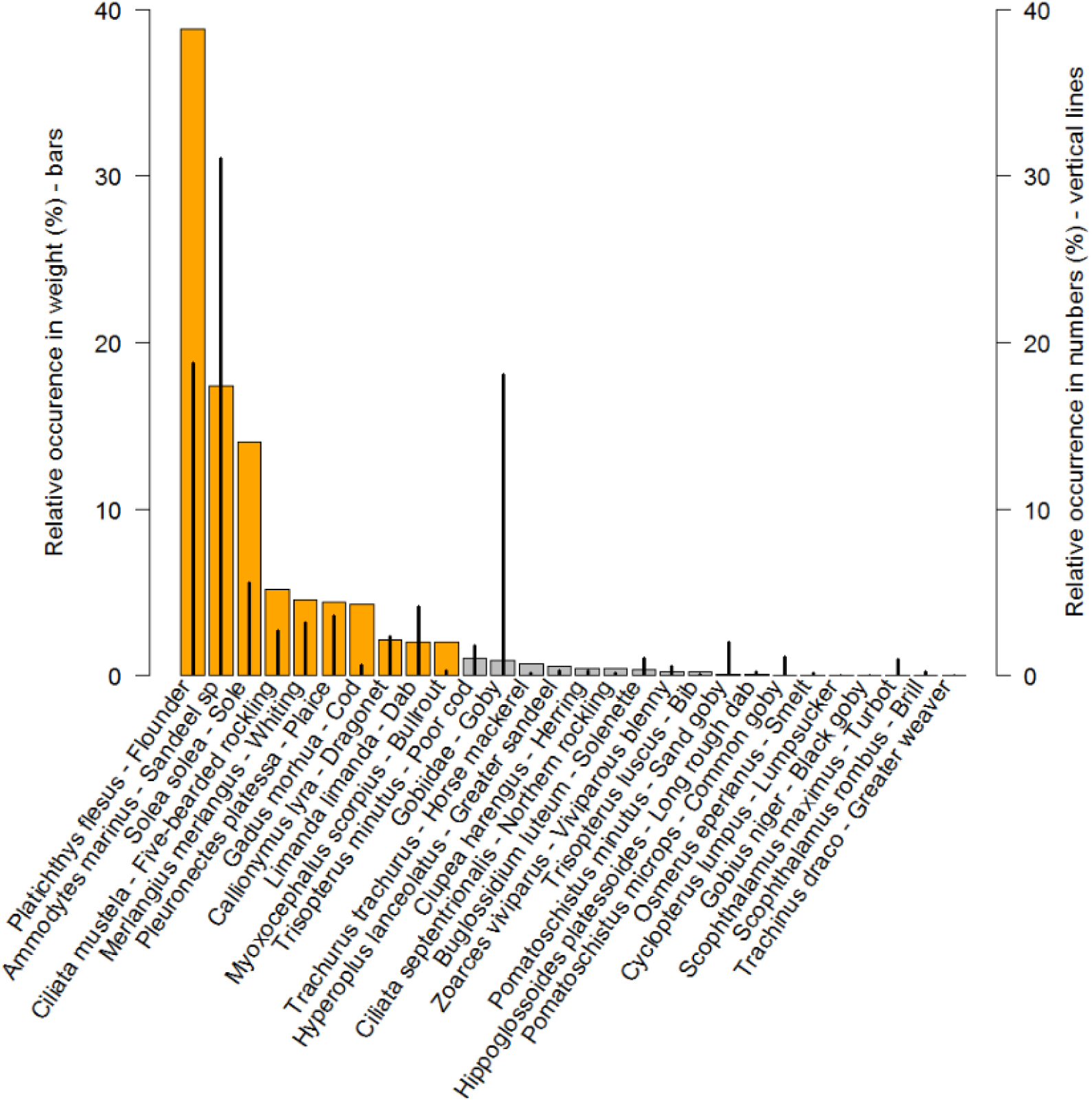
The percentage by estimated fresh weight (bars, left axis) and number of otoliths (vertical lines, right axis) of each fish species found in harbour seal scat samples collected in the Dutch Wadden Sea. The 10 most important prey species (95% of biomass in the diet) are indicated using orange vertical bars.

Estimated fish lengths in the faecal samples were mostly <25 cm, with a peak between 10 and 20 cm (Figure 4).

**Figure 4.**
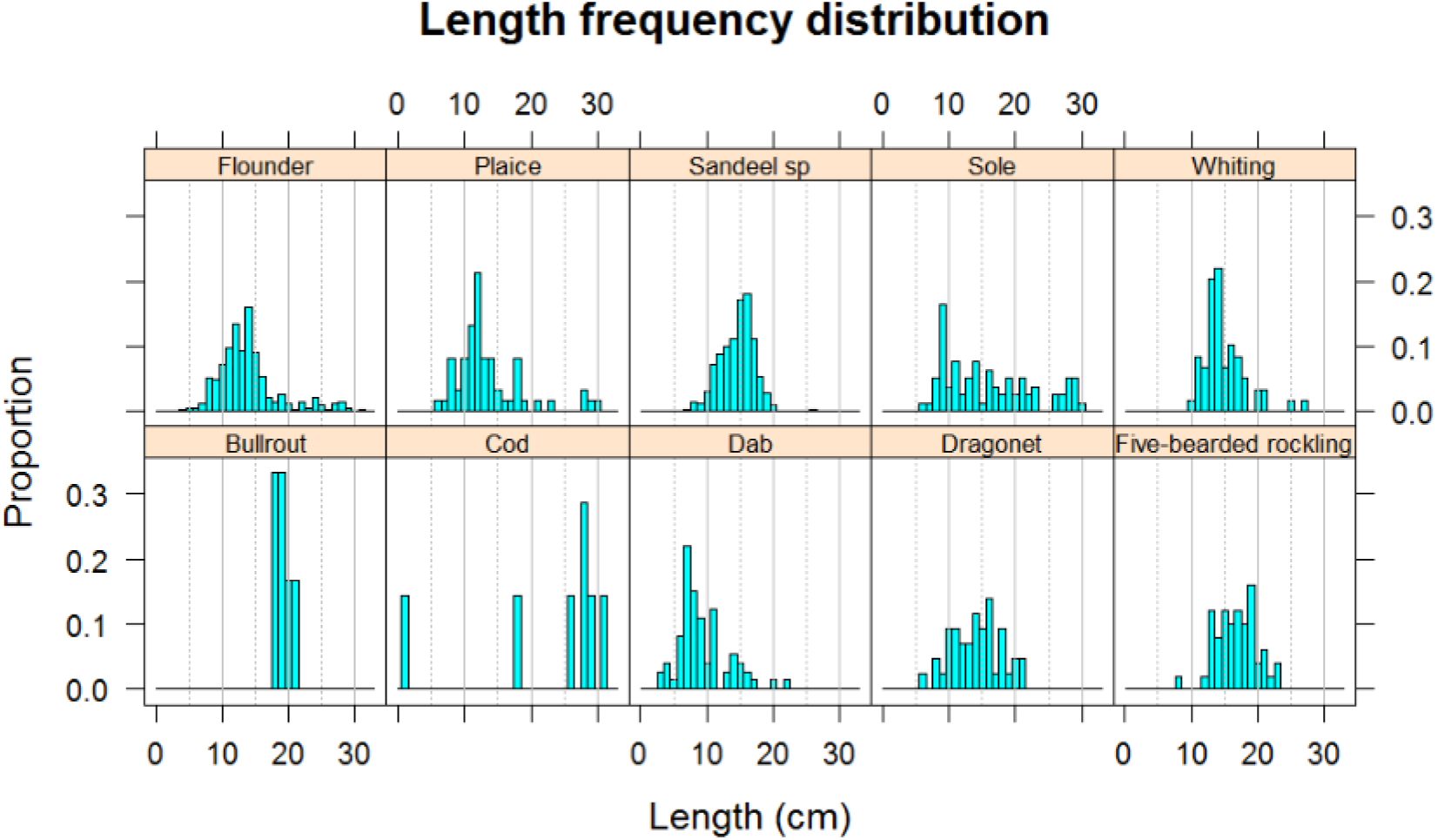
Length distribution of the 10 most common fish species found in scat samples of harbour seals, collected 1999-2009.

The estimated average energetic content of the fish consumed (weighted by their relative occurrence in the diet, see Figure 3) was 1.007 kcal/kg (Appendix S2, Table S2).

Assuming a daily energetic requirement of 4680 kcal per seal per day (Härkönen and Heide-Jørgensen 1991), this amounts to approximately 4.6 kg per day per individual.

#### Fish abundance

The biomass of the 10 most important species consumed by harbour seals in the Dutch Wadden Sea (i.e. plaice, sole, dab, flounder, sandeel, whiting, cod, five-bearded rockling, common dragonet and bull-rout) are shown in Figure 5 (and Appendix S2: Figure S4). Up to the mid 1980’s the fish biomass increased in both the Wadden Sea and adjacent coastal zone, but declined after that. In the most recent year (2016), the Wadden Sea contained on average 535 kg/km^2^ (corrected for catchability) and the adjacent coastal zone (Figure 2) held 2104 kg/km^2^. The total area of these areas are 2088 and 1707 km^2^, respectively. Hence, the average biomass observed in 2016 was 1117 tonnes for the Wadden Sea and 3592 tonnes for the adjacent coastal zone. In the Wadden Sea in the period between 2011 and 2015, the most abundant prey species were plaice (54%) and flounder (16%). In the coastal zone the most abundant prey species in the same period were dab (59%), whiting (28%) and dragonet (14%).

**Figure 5.**
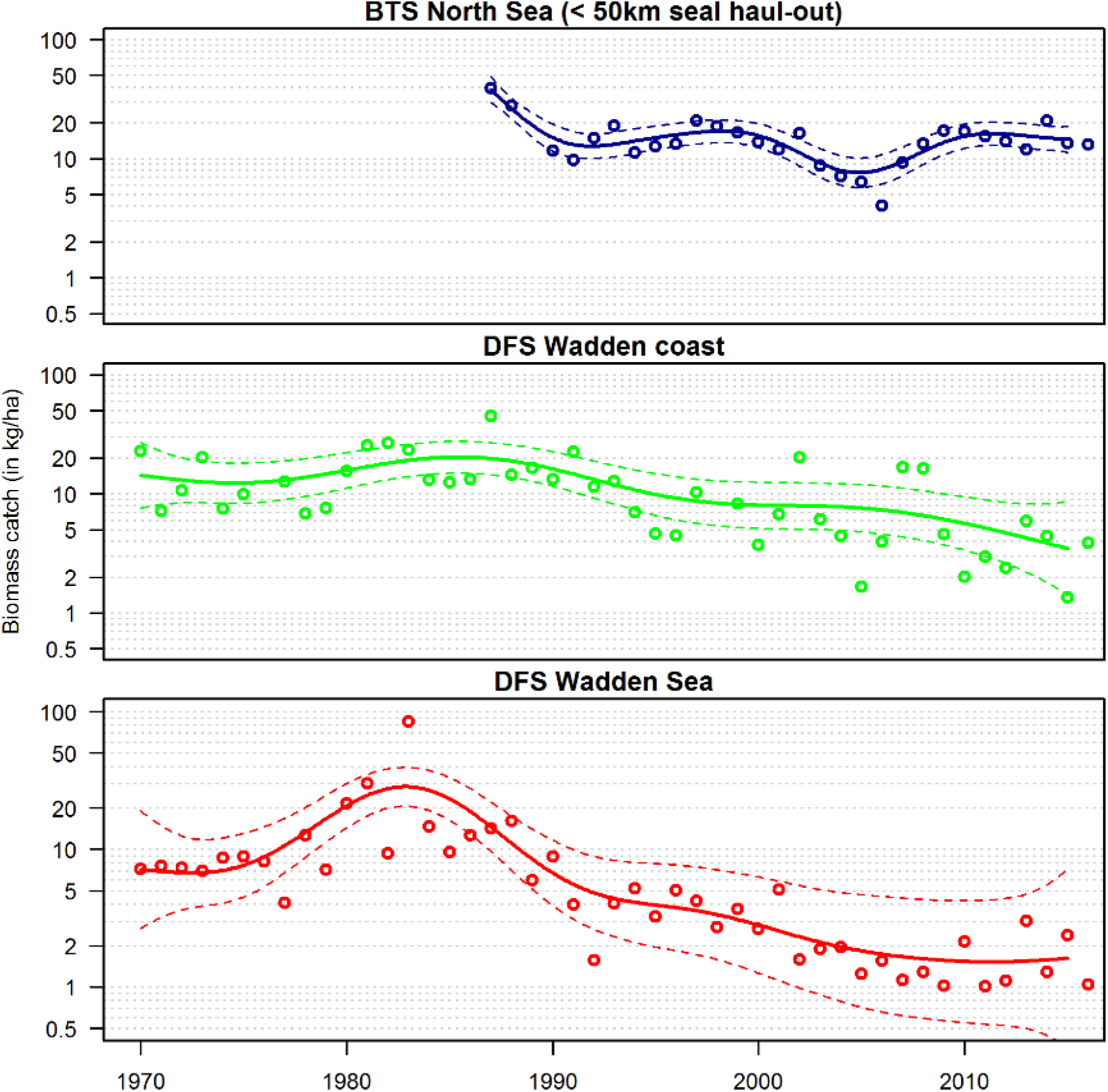
Trend in biomass (in kg/ha) of the 10 most important harbour seal prey species (based on scat samples), for the beam trawl survey (BTS) in the offshore zone, the demersal fish survey (DFS) in the coastal zone bordering the Wadden Sea (Wadden coast), and the DFS in the Wadden Sea (See also Figure 2). Note the log-scale on the y-axis.

#### Fish growth

The biomass of demersal fish in the Wadden Sea and nearby waters shows large seasonal variability. Most 0-year old fish (e.g. plaice, sole, and flounder) ‘settle’ in early spring (March/April). These 0-year olds can be highly abundant, but their total biomass is still low. They grow during spring and summer, and once they exceed ∼5 cm (around June-July, Figure 6, Appendix S2: Figure S3a-e) they can be caught by DFS-type gear. From September onwards, the apparent growth of 0-year old plaice and flounder is reduced substantially, despite the high water temperature (Figure 6). The following spring (March/April) they (now 1-year olds) start to grow, even though water temperature is still low (Figure 6, Appendix S2: Figure S3a-e). They continue to grow up to the late summer and fall. The predicted decline in length during the winter months is an ‘artefact’ of the DEB model: Since length is defined as a function of volume V^⅓^, a decrease in volume will lead to an apparent decrease in length.

**Figure 6.**
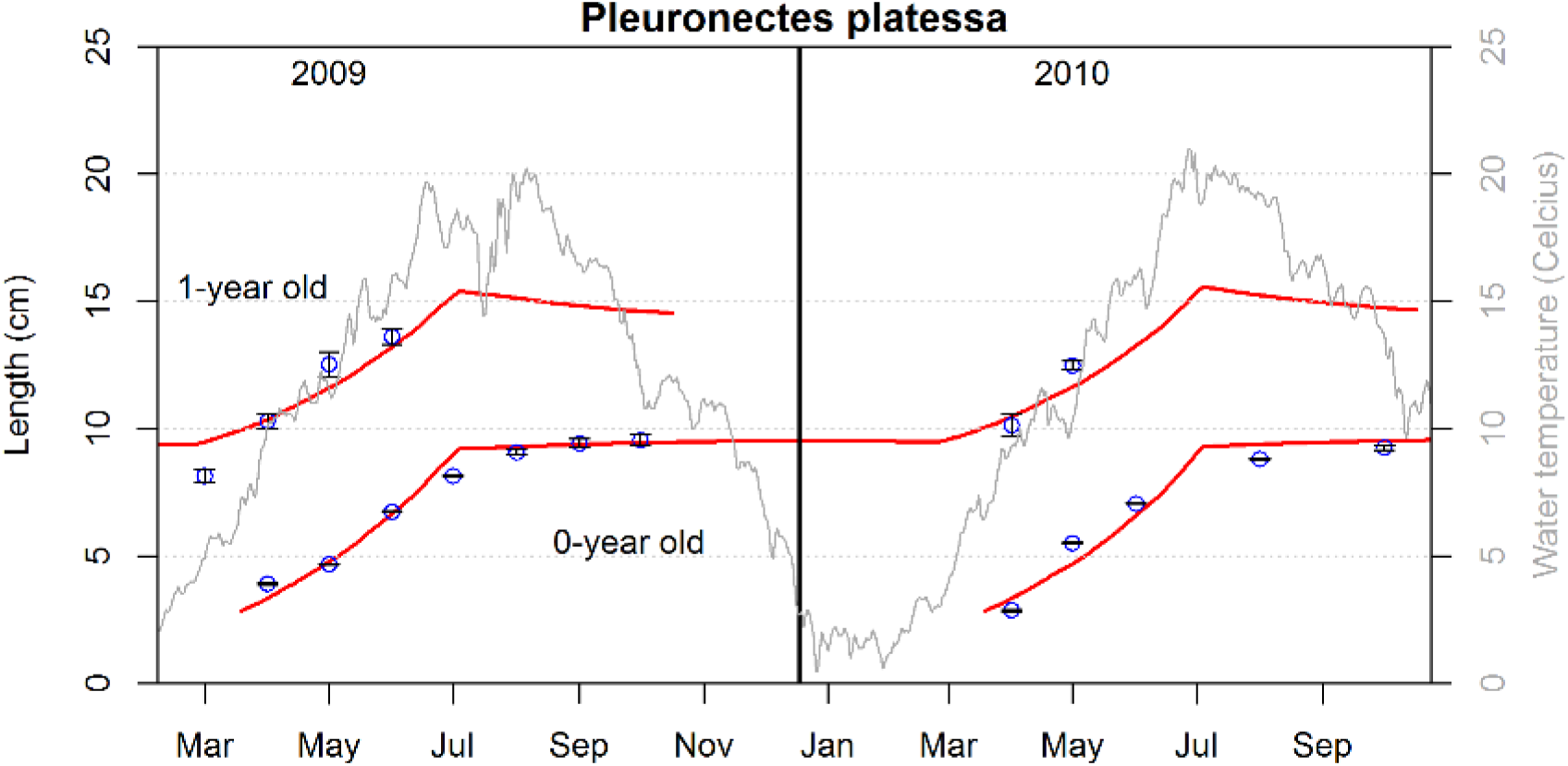
Observed length (circles and standard error bars) and predicted length (solid red line) of 0-and 1-year old plaice. Grey line indicates water temperature. Predicted growth (solid lines) is based on DEB model where food intake parameter *f* was allowed to vary between the winter months (*f*=0.11) and other times of the years (*f*=0.66).

#### Harbour seal spatial distribution and foraging activity

Although harbour seals are often observed on the sandbanks located within the Wadden Sea, on average they only spend 17% of their time on land (Appendix S2: Figure S1). Of the remaining 83% of their time, harbour seals spend on average 26% within the Wadden Sea, 28% in the adjacent coastal zone and the remaining time, 46%, in the offshore zone (Figure 2, Appendix S2: Figure S1). There are however, large seasonal variations. During summer months (April – September) seals spend most time on land (20-23%), and the least amount of time in the regions outside the 50km buffer. During the winter months seals spend most time further offshore beyond the 20m depth line (Appendix S2: Figure S1).

The time spent in each region is not necessarily a good proxy for foraging, as seals also perform other activities at-sea (like resting or transiting). When considering the total dive time (<1.5 m depth) in each region, only 14% of their dive time is spent within the Wadden Sea (substantially less than the total time spent there), 31% in the Wadden coast and harbour seals spend most dive time (46%) in the areas further offshore, particularly during the months January – March (63-74% of their dive time) (Appendix S2: Figure S1).

#### Consumption estimates

The total number of seals counted in the Dutch Wadden Sea in 2016, was 8160 (Galatius et al. 2017). Assuming a 0.68 haul-out probability during the Dutch aerial survey counts (Ries et al. 1998), 12,000 were estimated to be present. Based on the estimated 4.6 kg of fish consumed per seal per day, these 12,000 harbour seals in the Dutch Wadden Sea consumed approximately 56,000 kg of fish per day, and 20,400 tonnes of fish that year. In our study we excluded the Ems Estuary, and based our estimates on the remaining area, which contained 10,300 harbour seals 2016, consuming 17,500 tonnes of fish that year. The estimated consumption is substantially larger than the estimated fish biomass present in the Wadden Sea (ca 1100 tonnes) and adjacent coastal zone at the time of the survey (ca 3600 tonnes).

As fish growth is low during the winter months, harbour seals are expected to substantially reduce the total fish number in this period. This will affect the fish populations particularly when the estimated total numbers in September are low (e.g. 2011 in the Wadden sea; Figure 7). During the more productive spring and summer months, seals will continue to have an impact on the number of fish, however prey biomass continues to increase due to growth. The estimated consumption in the Wadden Sea and nearby coastal zone is presumed to be tightly linked to the number of fish present in the more offshore areas: when offshore fish density is high, the intake per unit of time (assuming a Type I functional response), will be higher (Figure 7). Figure 7 also shows the decline in numbers of fish (all ages) observed in a specific year, and the number of larger individuals (attempting to exclude 0-year olds) the following year. These calculations show that seal predation is likely a major contributor to the apparent annual fish mortality. However, for most years the fish mortality (or emigration) exceeds the estimated mortality caused by seals (the difference between solid coloured lines (with seal predation) and the dotted lines in Figure 7). This is particularly so for the offshore regions, but also for the Wadden Sea. For the Wadden coast the estimated impact is highest, and for several years the estimated mortality caused by seals exceeds the observed decline in prey numbers from one year to the next. Based on all years combined (2003-2016), the average estimated mortalities for the different regions are 43% (SD=13%) for the Wadden Sea, 60% (SD=19%) for the Wadden coast and 20% (SD=7%) for the North Sea region further offshore. Similar reductions were estimated for the biomass of the seals’ prey (Appendix S2: Figure S5). Since growth is limited during the winter months, seal predation was estimated to reduce the overall biomass during the winter months. However, during the productive summer months the increase in biomass strongly outpaced the seal predation.

**Figure 7.**
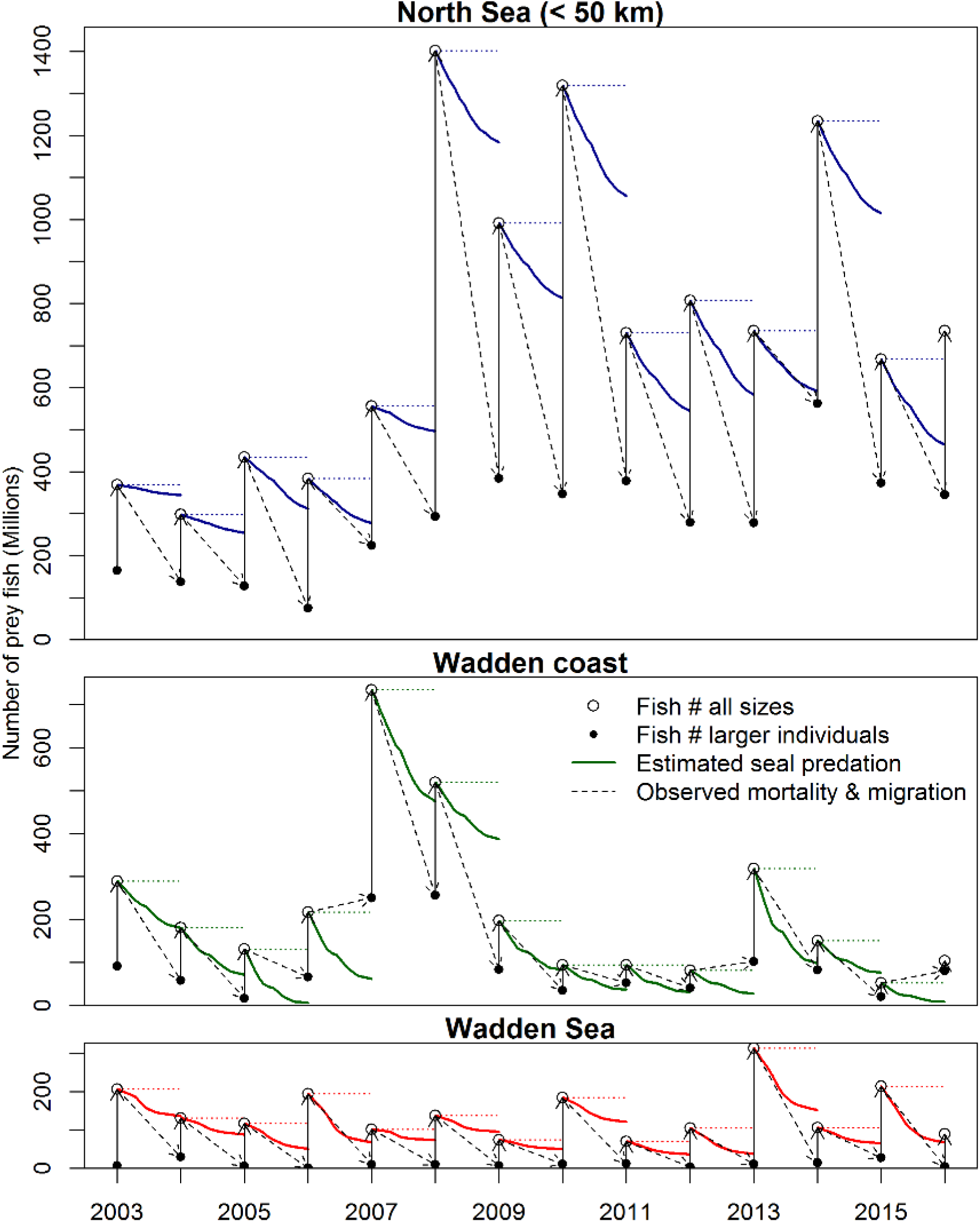
The estimated effect of seal predation on the number of prey fish for the Wadden Sea (red lines), Wadden coast (green lines) and remaining areas up to 50km from the haul-outs (blue lines, and see regions in Figure 2). The fish survey in September (i.e. BTS for the North Sea areas within 50 km of the nearest haul-out, and DFS for the Wadden coast and Wadden Sea) is the starting point (open circle, representing all size-classes), after which numbers decline due to seal predation. In the subsequent survey in September the following year, the number of remaining individuals (black dot, representing larger individuals, >13.5cm, presumed 1+ year olds) have declined, but is again supplemented with new recruits. In the Wadden Sea nearly all larger individuals (black dots) have died or disappeared (due to movement).

In the estimated impact of seal predation there were a large number of uncertainties. One of the largest source of uncertainty was the uncertainty in the catchability estimate, which is shown for one year (2016) in Figure 8. For the Wadden Sea, for 2016, the number of prey species still remaining in September (compared to the previous year) after seal predation was estimated to be 35% of what was present the preceding year (i.e. 65% reduction). However, the 95% CI was between 5% and 75%, implying that the estimated impact of seal predation could be both substantially larger or smaller than our mean estimate.

**Figure 8.**
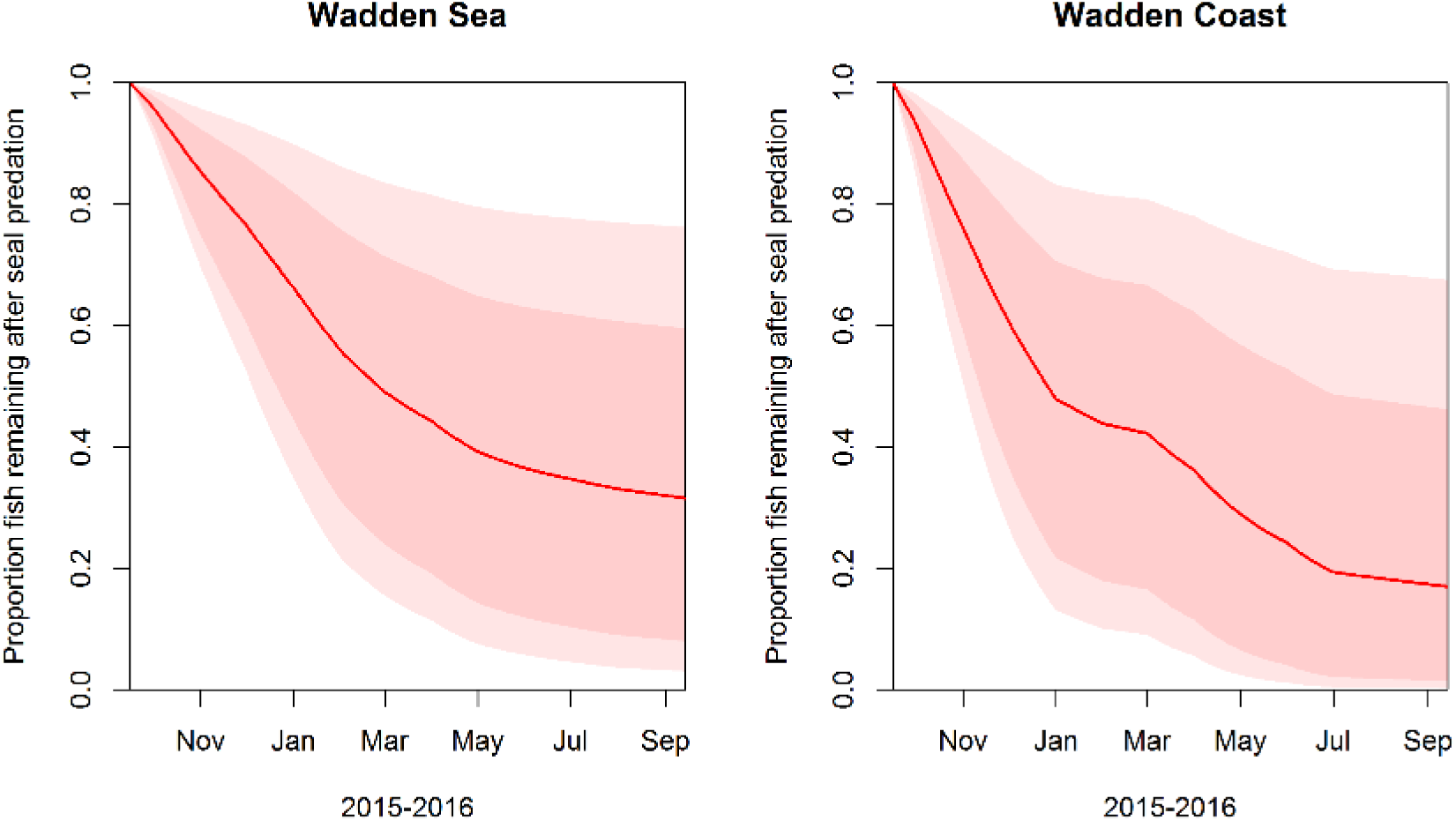
Uncertainty in the effect of the estimated impact of seals on fish numbers in the Wadden Sea (left) and Wadden coast (right). The solid red line represents the mean estimated impact, the darker pink area the ±SE and the light pink shaded area reflects the 95% confidence intervals.

## DISCUSSION

### The impact of seals on fish biomass in the Wadden Sea and adjacent coastal zone

We estimate that current harbour seal population imposes a considerable predation pressure on fish populations in the Wadden Sea and adjacent coastal zone. In our study area, the Dutch sector of the Wadden Sea (excluding the Ems estuary) and adjacent waters, the estimated annual fish consumption by harbour seals in 2015-2016 (i.e. 17 500 tonnes) was about three times higher than the combined prey fish biomass estimated for September in the Dutch Wadden Sea (1100 tonnes) and adjacent Wadden coastal zone (3600 tonnes). Since fish growth (i.e. biomass production) is small during the 5-6 winter months following the fish surveys, thus this region alone cannot sustain the entire harbour seal population.

This probably explains why harbour seals only spend ∼13% of their diving time in the Wadden Sea. We estimated that seals consume approximately half of the fish present in the Wadden Sea from September onwards. In the coastal zone bordering the Wadden Sea, we estimated that the seals exert a similar predation pressure. There, the total fish biomass is higher than in the Wadden Sea (particularly in recent years), but seals also spend more time foraging in this region.

Although the total demersal fish biomass in the Wadden Sea and nearby waters has decreased in the last two decades and the size of the harbour seal population has strongly increased, it is unlikely that seals are the single cause of the decline in the fish populations in the Wadden Sea. The decline in fish stocks in the Wadden Sea and coastal waters started in the mid-80’s, when seal numbers were still around their historic low. Most probably the decrease in fish biomass in the mid-1980s is related to the decrease in eutrophication and climate change (Teal et al. 2012, Støttrup et al. 2017). Furthermore, the decline in demersal fish biomass (and fish numbers) in the coastal shallow waters occurred in all age-classes including the 0-year olds (Støttrup et al. 2017). The majority of fish in the seals’ diet in our study area was larger than 10cm (Figure 4). Most 0-year olds only reach that size at the end of the growth season (Appendix S2: Figure S3). Since we expect the predation pressure by seals on these 0-year olds prior to the survey in September to be rather low, we assume that bottom-up processes (e.g. low nutrient loadings) in combination with mortality by birds or by-catch in the shrimp fishery is a more likely explanation for the observed decline in 0-year old fish biomass. However, during the winter months following the survey, seals can impose a considerable predation pressure on the remaining prey. Hence, given the low fish density and high seal numbers it is likely that at their current numbers seals are applying top-down pressure on local fish populations, especially with regard to preferred prey species.

These results are in line with some studies that suggest that the predation pressure by marine mammals can alter the abundances of prey species. For examples, grey seals (*Halichoerus grypus*) may have impaired the recovery of over-exploited north-west Atlantic cod stocks (Trzcinski et al. 2006, Cook and Trijoulet 2016). Benoît et al. (2011) estimated that in that region, grey seal predation could explain 20-50% of natural mortality of some demersal fish species, even if these species comprised a small percentage (<25%) of the grey seal diet. In contrast, other studies elsewhere have demonstrated low impacts by seals on fish communities (Houle et al. 2016). Such differences between observed impacts undoubtedly relate to the sizes of seal and fish stocks, the relative importance of other ecosystem components (e.g. predatory fish like cod and whiting, Temming & Hufnagl 2015), and stock replenishment rates. The Wadden Sea could be considered the seals’ back-garden. In the vicinity of the haul-out sites from which they forage seal predation is most likely to cause food depletion, a principle known as Ashmole’s halo (Gaston et al. 2007).

### Sources of uncertainty

One main source of uncertainty was the lack of experimentally derived catchability estimates for the DFS gear. Instead, we attempted to estimate catchability based on experiments for plaice, which used different fishing gear (Kuipers (1975)). The catchability of some species might be very different from plaice. For example sandeel is the second most important prey item observed in the seal diet, but they are not caught optimally by the used gears because of their pelagic and burial phase, and their thin elongated shape. Surveys specifically designed for sandeel use gear that dislodges them from the substrate notably catch higher numbers (Tien et al. 2017). Also other pelagic species (e.g. herring, sprat, etc.) are poorly sampled by a demersal beam trawl gear (Couperus et al. 2016). Although only two of the sampled scats contained herring (max 5%), only five samples were collected between December-February. Harbour seals may feed more on pelagic fish species (e.g. herring, sprat, etc.) during these winter months (de la Vega et al. 2016). This could imply that the consumption of demersal prey, and impact of seals on the demersal fish community, might be overestimated.

Another unknown factor, is where seals capture their prey. In this study, the prey removal in each region was defined as the product between prey density and dive time in the different regions. Clearly, some dive time is not related to foraging, and the amount of non-foraging dives may differ between the regions, and even between seasons. Also the distribution of their prey is not static. While, we accounted for changes in prey density due to seal predation, relative changes in fish distribution were not taken into account. For example, several studies have suggested an offshore movement of fish toward deeper (warmer) waters during the winter months (de Veen 1978, Teal et al. 2012, van der Veer et al. 2016). This also seems to be reflected in the movement of seals, who tend to forage further offshore during the winter (Aarts et al. 2016). In addition, also other sources of mortality (e.g. predation by cormorants or bycatch in the shrimp fishery (Leopold et al. 1998, Glorius et al. 2015)) could locally lower fish density. This would lead to lower intake rates by seals, possibly over-estimating the effect of seal predation in the Wadden Sea.

However, regardless of where exactly seals capture their prey, all living seals ultimately need to acquire sufficient prey to meet the energy demands (Härkönen and Heide-Jørgensen 1991). Since foraging sites within and close to the Wadden Sea require less travelling, they are more beneficial. Therefore optimal foraging theory would suggest the highest reduction in food density to occur near the colony.

### Other piscivorous ‘predators’ in the Wadden Sea

In order to evaluate the relative importance of top-down regulation by harbour seals, we need to consider other sources of mortality, such as predation by other marine mammals, piscivorous birds, predatory fish, but also commercial fishery (Zijlstra and Van Eerden 1995, Leopold et al. 1998, Arnett and Whelan 2001, Glorius et al. 2015, van Kooten et al. 2015, Hansson et al. 2017).

Grey seal numbers in the Wadden Sea grew exponentially after 1990 (Brasseur et al. 2015), with a maximum of 4045 counted in the Dutch section of the Wadden Sea during the moult in 2017 (Brasseur et al. 2017). Although grey seals in the North Sea primarily feed on sandeel, they also feed on other benthic prey species (Brown et al. 2012). While grey seals may visit areas >100km offshore, presumably to feed on sandeel banks, they also spend a large amount of time in coastal waters near the Wadden Sea, overlapping with harbour seal distribution.

In addition to grey and harbour seals, 30-80 thousand harbour porpoises are estimated to reside in the Dutch section of the North Sea (Geelhoed et al. 2013). Only a small part of this population uses the Wadden Sea and nearby coastal zone. In Denmark and Germany harbour porpoises are present near the Wadden Sea throughout the summer season, and are known to feed on 0-year old flatfish species (Gilles 2008). However, stomachs of harbour porpoises found stranded along the Dutch coast, mainly contain whiting, sandeel and gobies and only a small proportion of flatfish (Leopold 2015).

Several bird groups e.g. cormorants and divers also feed on demersal fish species. Cormorants are abundant and specialize mainly on 0-group flatfish, e.g. plaice, dab and flounder (Leopold et al. 1998). Currently approximately 25,000 cormorants breed in the Netherlands, of which half live in the vicinity of the Wadden Sea (www.sovon.nl). Cormorants require approximately 460 – 500 g of fish per day (Zijlstra and Van Eerden 1995, Leopold et al. 1998). Assuming these cormorants feed continuously within and near the Wadden Sea, this equates to a prey requirement of approximately 2300 tonnes per year.

In addition to the natural sources of mortality, mortality due to commercial fisheries should also be considered. In the North Sea, the total catch by the commercial fishery exceeds by far the consumption by marine top-predators (Engelhard et al. 2013). A pattern which is also evident in other regions, like the Baltic (Hansson et al. 2017). However, locally near the coast with high densities of central-place foraging predators, the impact of these predators may exceed that of fishing (see e.g. also (Hansson et al. 2017)). In the Dutch coastal zone, the fisheries targeting demersal fish species is rather small (i.e. landing 723 tonnes, including 99 tonnes in the Wadden Sea (van Kooten et al. 2015)). The shrimp fishery is by far the most important fishery in and around the Wadden Sea, with plaice being the most important by-caught species. In de Dutch Wadden Sea alone, shrimp fisheries catch an estimated 99 million 0 and 1-year old plaice, mostly during the summer months (adapted from (Glorius et al. 2015)). The total number of plaice (corrected for catchability) present during the DFS in September was estimated at 89 million. This suggests that prior to the DFS, approximately 50% of plaice was already caught away by the shrimp fishery.

Although the population size, food requirement and target species differ between marine predators and fishery, they can collectively impose a considerable predation pressure on the demersal fish communities of the Wadden Sea and nearby coastal waters. Recent simulations for a North Sea ecosystem not only suggest that top-down fishing pressure can be of tremendous importance for the dynamics of fish populations, the top-down effects may also cascade down to lower tropic levels, including plankton abundance (Lynam et al. 2017). This might also be the case for the effect of predators, such as seals, in coastal areas.

### Are harbour seals food-limited?

The estimated food requirement by harbour seals is well above the amount of prey estimated to be available in the Wadden Sea and Wadden coast combined. Since harbour seals are central-place foragers, some depletion near the colony is likely to occur. However, seals also forage well beyond the coastal zone (Figure 2), where prey density is higher (Figure 5) and the estimated depletion is substantially lower (Figure 7). In recent years, harbour seals appear to be using the west coast of the Netherlands more frequently (Aarts et al. 2013), which suggests that more distant areas have become more attractive foraging locations than previously. Although the harbour seal population in the Dutch part of the Wadden Sea is still increasing (but at a slower rate than before), in other regions, like Denmark, the population seems to have reached a plateau (Brasseur et al. 2018). Given the estimated impact of harbour seals on the fish community, it is likely that the slowing down in the population growth is at least partly the result of food limitation.

### Density dependent processes and possible interactions between seals and fishery

While seals can impose a considerable mortality on individual fish, it is still debatable whether seals have an overall impact on fish biomass. This ultimately depends on whether seal predation keeps the fish numbers well below the level where intra-or inter-specific competition for food will occur. Lorenzen and Enberg (2002) reported density dependent growth (DD-growth) reduction in 9 out of 16 marine fish populations studied. DD-growth is assumed to mainly occur in the early life of a cohort (Anderson et al. 2017) and has been reported frequently in species that concentrate during their juvenile phase in shallow coastal waters (Beverton 1995). For instance, DD-growth has been observed in juvenile plaice and sole in years of exceptionally large year classes (Rijnsdorp and Van Leeuwen 1996). The observed reduction in growth of 0-group fish species (e.g. plaice and flounder) corroborates that in summer food becomes a limiting factor in the Wadden Sea.

The density-dependent reduction in juvenile growth may have been a regular phenomenon prior to the period of eutrophication of the coastal waters (Bolle et al., 2004). Productivity in the coastal waters has decreased recently supposedly in response to the decrease in nutrient inputs (Philippart et al. 2007, Støttrup et al. 2017), lowering the biomass at which density-dependent competition for food will occur. Whether the increased predation pressure from seals is sufficient to suppress the fish biomass below a critical level and limit where resource competition starts to slow down growth requires further study.

Another aspect that would require further study, is whether seals compete with other predators, like the commercial fishery. When both marine mammals and fishery target the same (over-exploited) fish species, the additional mortality imposed by seal predation may hamper the recovery of some commercially important target species, as was suggested for cod in the west Atlantic (e.g. Trzcinski et al. 2006, Cook et al. 2015, Cook and Trijoulet 2016). However, if marine mammals and fishery target different species or size classes, this could give rise to both positive and negative feedback loops, and the competition between marine mammals and fishery might be less obvious (Morissette et al. 2012, Houle et al. 2016).

A number of harbour seal prey species, e.g. sole and plaice, are also targeted by the commercial fishery in the Southern North Sea. The Wadden Sea and adjacent coastal areas are important nursery grounds for these species (van der Veer et al. 2011). Seals feeding in these areas target fish before they move further offshore, and become available to the fishery. This leads to the question whether or not the seal predation could diminished catch opportunities for fisheries on these species later on. This depends on how these populations are regulated (De Roos et al. 2007). When density-dependent growth occurs in the juvenile stage, increased mortality can actually lead to higher biomass of adults, because it removes the bottleneck which limits recruitment into larger age-classes (van Kooten et al. 2007, De Roos et al. 2007). Seal predation would only translate into a proportional reduction in stock size for fisheries when seal mortality linearly translates into reduced numbers of adult fish, with no effects on growth. This would be the case if seals would feed on the same sized fish that the fishery targets. Interestingly, we find that the biomass of the species caught in the BTS (aimed at sampling the larger commercially important flatfish species), in the region within 50 km of seal haul-out sites has increased slightly in recent years (Figure 5), a pattern also observed elsewhere (Støttrup et al. 2017). However, this is not necessarily a positive effect of seal predation (for example, beam trawl fishing intensity has also strongly diminished in the area), at least the increased seal presence does not coincide with a decrease in fish abundance further offshore.

## ACKNOWLEDGEMENTS

We want to thank all crew and assistants from Wageningen Marine Research that assisted in the field with the seal work and collected the fish data for all those years and especially: Piet-Wim van Leeuwen, Andre Meijboom, Hans Verdaat, Marcel de Vries, Andre Dijkman, Gerrit Rink & Thomas Pasterkamp and the crew of the Wadden Unit. The Dutch ministry of Agriculture, Nature and Food Quality (MinLNV), Groningen Sea Port, Eneco and Gemini windpark funded the seal GPS transmitters. MinLNV funded the aerial surveys in the Wadden Sea and we thank the pilots, in particular Aad Droge. The National Ocean and Coastal Research Programme survey was financially supported by the Netherlands Organization for Scientific Research (NWO). The DFS survey was carried out as part of the statutory tasks set out in Dutch legislation on fisheries management, financed by MinLNV. This research was partly funded by the MinLNV, and the study was partly funded by the “KennisBasis” program System Earth Management internally lead by Martin Baptist, project number KB-24-002-020. Finally, we thank Sophie Smout for providing usefull comments on the manuscript.

To enter protected areas and handle seals during field procedures, the following permits were obtained: A permit under the Dutch Nature Protection Act (Natuurbeschermingswet) given by the Province of Friesland, a permit under the Flora and Fauna Act (Flora en Fauna Wet) given by the Dutch government and protocols approved by an animal ethics committee (Dier Ethische Commissie, DEC) of the Royal Netherlands Academy of Science (KNAW).

## APPENDIX S1. Estimating catchability of the fish surveys

Estimating the catchability of the survey gears is essential for estimating absolute fish abundance, and hence for estimating the impact of seals on fish stocks. Two approaches exist for estimating catchability. The first approach is to estimate the total stock size using stock assessments, and subsequently comparing the catch per unit of area with the expected numbers per unit of area based on these assessments (e.g. (Fraser et al. 2007, Walker et al. 2017)). Because the survival rates for the younger age classes (e.g. 0 and 1-year olds) are not well known the stocks size estimates of these age-classes are likely inaccurate. As a result the derived catchability estimates for these age-classes will be poor.

The second approach to estimate catchability involves experimentally estimating the proportion of fish escaping the net. (Kuipers 1975) defines four main processes that cause fish to escape the net: 1) Lateral escape, 2) Disturbance caused by the ship, 3) Escape through the net and 4) Escape underneath and above the net. (Kuipers 1975) attempted to experimentally estimate the effect of these 4 processes using a 1.9m wide small meshed beam trawl. This resulted in a catchability equation which was specified as function of the beam width. Here we re-estimated that function, and use it to estimate the catchability of the 3m and 6m DFS gear.

### The efficiency of a 1.9 m beam trawl

The experimental study area was located on the tidal flats of the Balgzand, in the Dutch Wadden Sea. Fishing was carried out by a rubber dingy, fishing at a speed of 0.5-0.58 m/s, and a stretched mesh size of 1cm, using 1.9 m (or 3m) beam and 0-3 tickler chains. Given the relative small (stretched) mesh size of 1cm, only fish <2.5 cm were considered to partly disappear through the net (item 3 above). Catch rates depended on whether tickler chains were used, but fishing with more than one tickler chain did not lead to higher observed catch rates. Hence, the proportion of fish escaping underneath the net was assumed to be negligible when fishing with more than 1 tickler chain (item 4 above).

To estimate the effect of **boat disturbance**, catch rates were compared when fishing with a 8m and 400m long fishing line. The observed discrepancy between catch rates increased with length. The efficiency E_d_ as function of length was provided, but this did not match the function provided in the corresponding figure of (Kuipers 1975). Hence, the data were reconstructed (Figure S1) and a new efficiency function was estimated.

**Figure S1.**
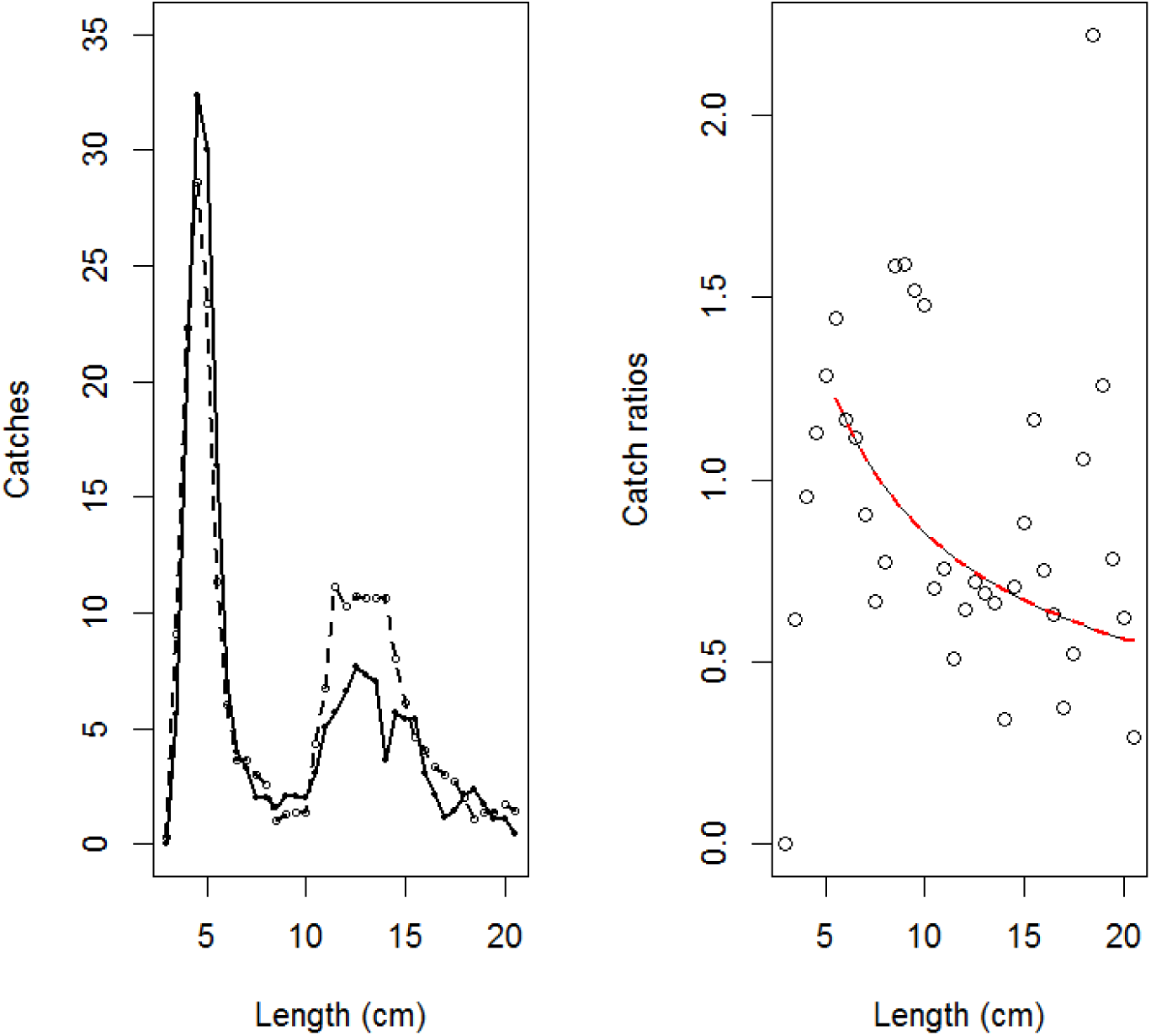
(Figure 5 in (Kuipers 1975)). Catches of plaice by length class of the 2m beam trawl with 8m (solid line) and with 400m (dashed line) fishing line (moving averages of 3). Right, efficiency in respect of disturbance (E_d_) against plaice length. Curve represents the log of catch-ratio as function of log of length.

A Generalized Linear Model (GLM) assuming a quasi-Poisson distribution was fitted to the data. The response variable was defined as the catch rate of the 8m fishing line, and modelled as a function of the log of fish length (*L*), with the log of the catch rate of the 400m fishing line included as an offset. Similar to (Kuipers 1975), only data from fish > 5cm were included in the model fitting. This results in the following efficiency (as fraction) caused by boat disturbance

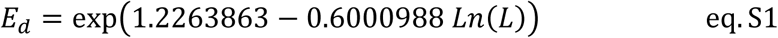

The effect of **the lateral escape** was estimated by comparing the catch rates of a 1.9m beam and two 1.9m beam fitted together side-by-side. Similar to the analysis based on the 8 and 400m fishing line, a GLM was fitted to the data, with the catch rate of the single beam as the response variable, and the log of the catch rate of the double beam (i.e. 2 × 1.9m) as model offset (Figure S2)

**Figure S2.**
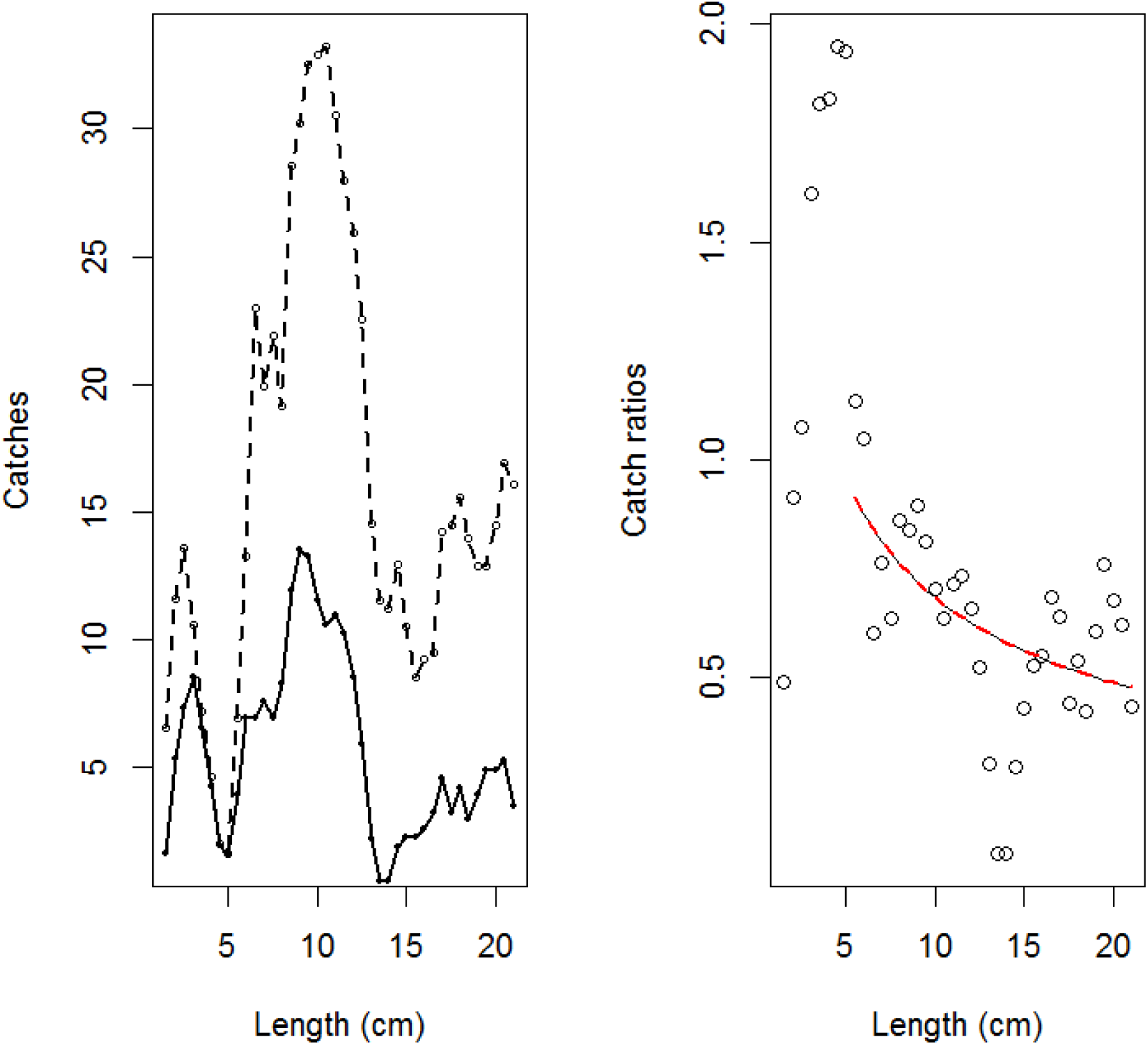
(Figure 6 in (Kuipers 1975)). Catches of plaice by length class of the 1.9m beam trawl (solid line) and 2×1.9m beam trawl (dashed line) (moving averages of 3). Right, catch-ratio in respect of lateral escape (E_l_) against length of the plaice. Note that the catch rates (right figure) were corrected for beam-width. Curve represents the log of catch-ratio as function of log of length.

This resulted in the following catch-rate ratio of the 1.9 m beam (*c*) relative to the catch rate of the 2 × 1.9m beam (*C*).

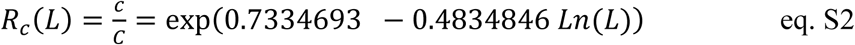

The philosophy underlying the lateral escape, is that within a fixed section (width = *b* cm) on either side of the net, fish escape. Hence, the ratio between the catch rate can be defined as

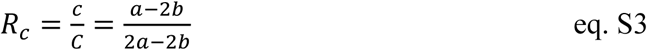

Where *a* is the size of the single net (i.e. 1.9m) and 2a the size of the double net. The size of *b* is length dependent, with *b* being larger for larger fish, due to their higher swim speed. Hence, *b* can be estimated as

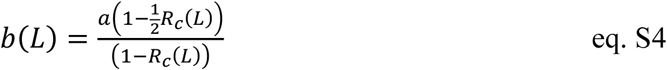

The total efficiency regarding the lateral escape of the 1.9 m relative to no lateral escape can now be defined as

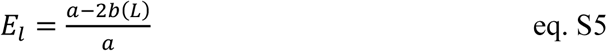

, where here *a*=1.9m.

The total efficiency *E*, taking both lateral escape and boat disturbance into account, can be defined as

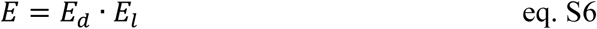

### The efficiency of the DFS beam trawl

Similar to the gear used by (Kuipers 1975), the DFS uses a net with a similar mesh size (i.e. 2cm) and 1 tickler chain. However, the DFS differs in a number of other aspects. 1. The survey takes place at a higher speed (i.e. 1.03-1.54 m/s, instead of 0.5 m/s). 2. It uses a larger beam width (3 m in the Wadden Sea, and 6 m in the Wadden coastal zone), 3. Most fishing takes place in the Wadden Sea gullies, and not on the relative flat tidal mud flats as in, which may lead to a different proportion of escape underneath.

Using eq. 5 it is now possible to correct the efficiency to take the larger beam width into account (Figure S3). (Kuipers 1975) did not find evidence for fish escaping underneath the net when using at least one tickler chain, but fishing took place on the relative flat mud flats. In contrast, (Reiss et al. 2006) towed three nets behind each other, and found that on average only 37% of the flatfish dab (*Limanda limanda*) remained in the first net, while the others escape underneath. This might be due to a difference in behaviour between dab and plaice, but could also be due to a difference in topography and bottom structure. Since the DFS surveys the gullies and coastal waters of the North Sea, we assume that the proportion of escape underneath is better represented by the study of (Reiss et al. 2006). When combining the reconstructed length-specific efficiency and accounting for possible escape underneath the net based on (Reiss et al. 2006), the efficiency becomes substantially lower (solid red and purple line, Figure S3). These efficiency estimates will be used in the subsequent calculations.

We expect this to be an under-estimate of efficiency, particularly because the DFS fishes at twice the speed of the (Kuipers 1975) study. However, there appears to be a close correspondence with the efficiency estimates from (Bergman et al. 1989) (green line and solid purple line in Figure S3), who also used a 3m beam trawl. However, unfortunately no details or data are presented in (Bergman et al. 1989), and hence their estimated efficiency cannot be validated.

**Figure S3.**
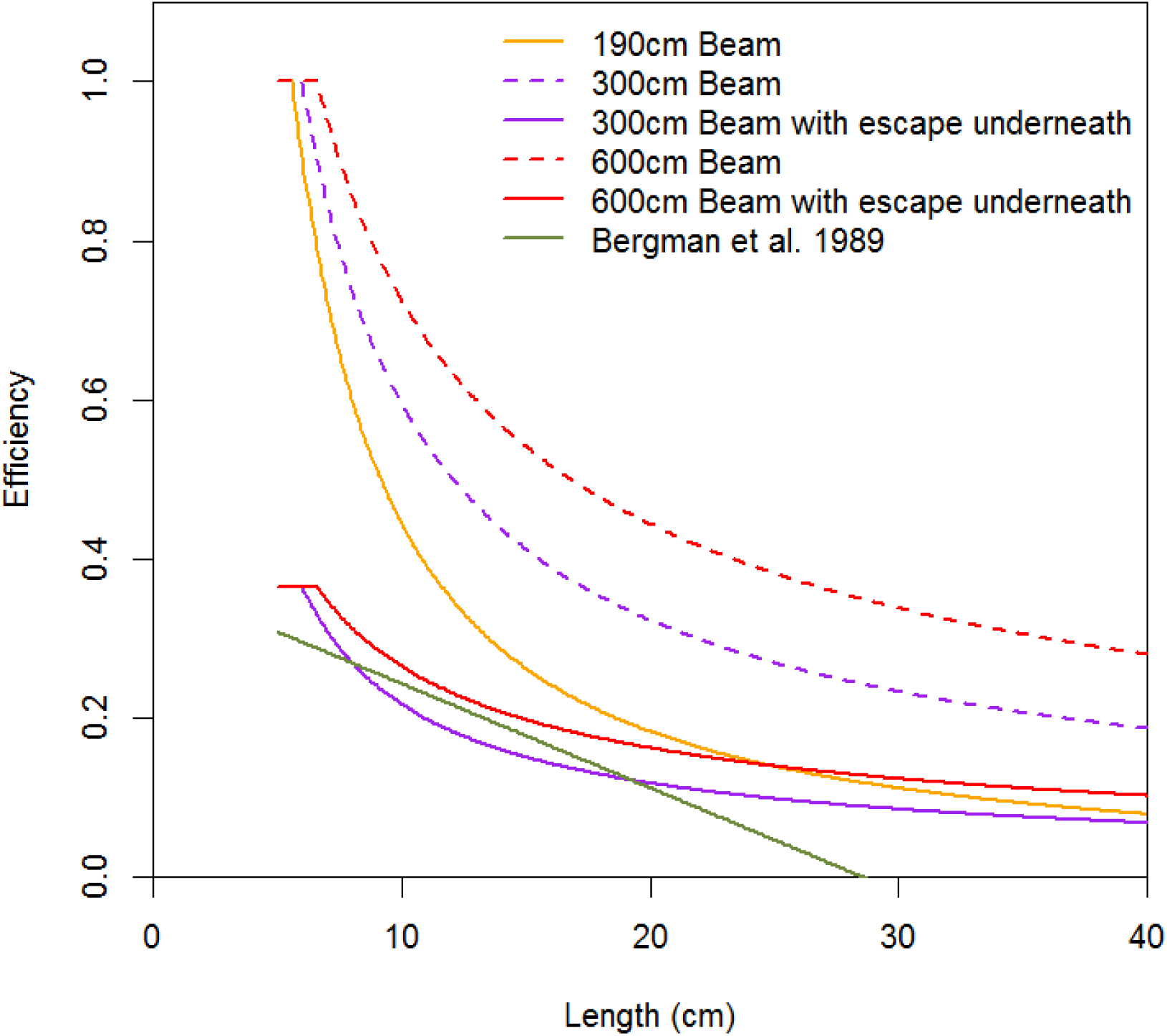
Efficiency estimates for flatfish for different gear types taking the size-selective catchability into account. The orange line is the total estimated efficiency for plaice based on (Kuipers 1975). The dashed purple and red line are based on (Kuipers 1975), but correcting for the larger beam-width (i.e. 3 and 6m, respectively). The solid purple and red line, also take escape underneath into account (Reiss et al. 2006). These lines will be used as the catchability estimates of the DFS in the Wadden Sea (3m beam) and Wadden coastal zone (6m beam). The green line represents the estimated efficiency of a 3m beam based on simultaneously fishing with a 1.9 and 3m beam trawl (Bergman et al. 1989). Catchability estimates exceeding 1, where truncated to 1.

### The efficiency of the IBTS, SNS and BTS

For the regions bordering the Wadden Sea, data from the international bottom trawl survey (IBTS), sole net survey (SNS) and beam trawl survey (BTS) are available. Although, surveys also take place in a larger area, further offshore, some fish hauls occur in the same region as the coastal DFS. For those sampling stations in the overlapping years (here data from 1987 onwards were used) it was possible to directly compare the length-specific catch per unit area estimates of the IBTS, SNS and BTS survey, with those from the DFS. This will provide a measure of relative efficiency. Since, we now have a (rough) estimate of the efficiency of the DFS, we can reconstruct the absolute catchability.

95% of the DFS sampling locations occur at depths shallower than 20m, hence we only select the IBTS, SNS and BTS survey data from similar depths and within at least 50km of the nearest seal haul-out site in the Wadden Sea. The catch-ratios are shown in the top-figures of Figure S4a-c. Up to approximately 10 cm, the catch per unit of area are higher in the DFS, compared to the BTS and SNS, which is possible due to the larger mesh size of the latter two gears (i.e. 4 cm). However, catch-rates are substantially larger at larger length classes. For example, for a length-class of >20cm, the BTS and SNS catch at least 3 times more individuals per unit of area. This is most likely due to the larger fishing speed (BTS/SNS = 4nm, DFS= 2-3 nm). Overall, the catch rate of the IBTS is much lower, which is most likely due to the fact it does not use a heavy beam.

**Figure S4a.**
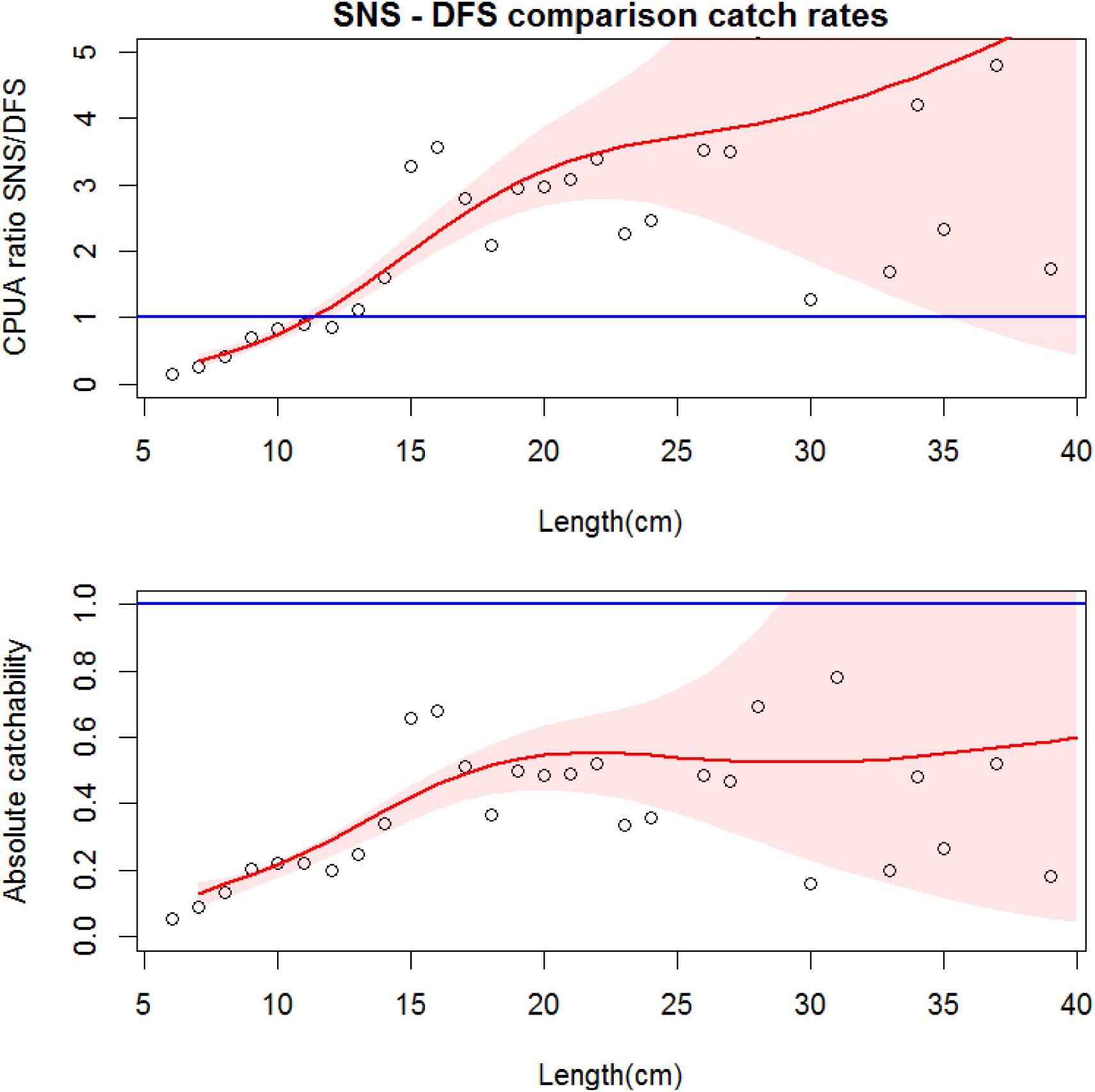
Catch ratios for plaice of SNS and DFS (top figure). Values <1 imply that the absolute numbers per ha caught in the DFS are higher than in the SNS. Red line is a GAM fitted to the data (shaded area is 95% CI). Lower figure is absolute catchability of the SNS survey, taking the catch-ratio between SNS/DFS and the absolute catchability estimate of the DFS (Figure S3, red solid line) into account.

**Figure S4b.**
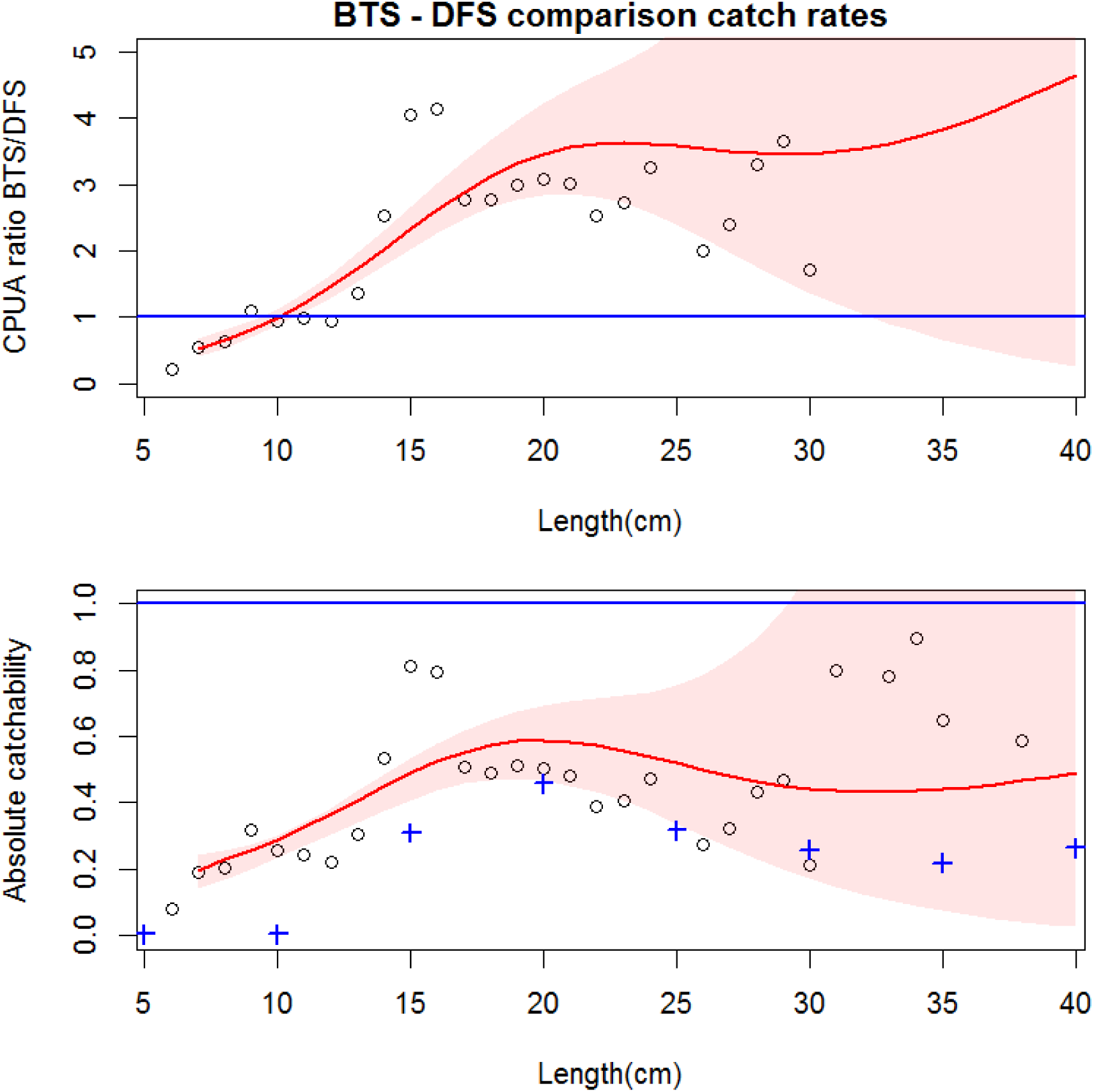
BTS catchability estimates. See Figure S4a. Blue plusses are the absolute catchability estimates for plaice based on (Walker et al. 2017). The higher catchability estimates for the larger length classes, might be explained by a higher turbidity in the coastal waters, possible causing lower lateral escape.

**Figure S4c.**
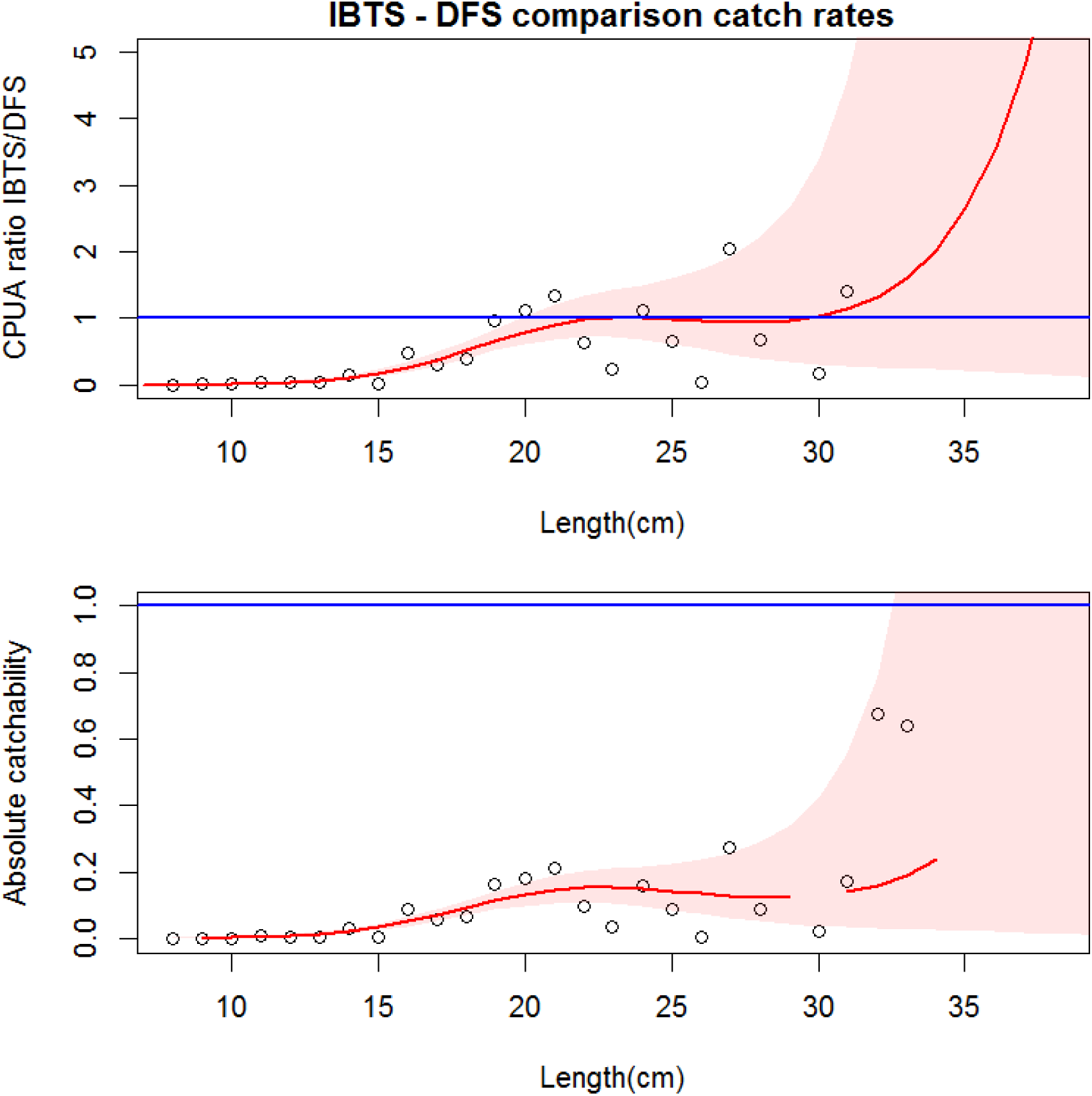
IBTS catchability estimates. See description Figure S4a.

## APPENDIX S2. Supplementary figures and tables

**Table S1.**
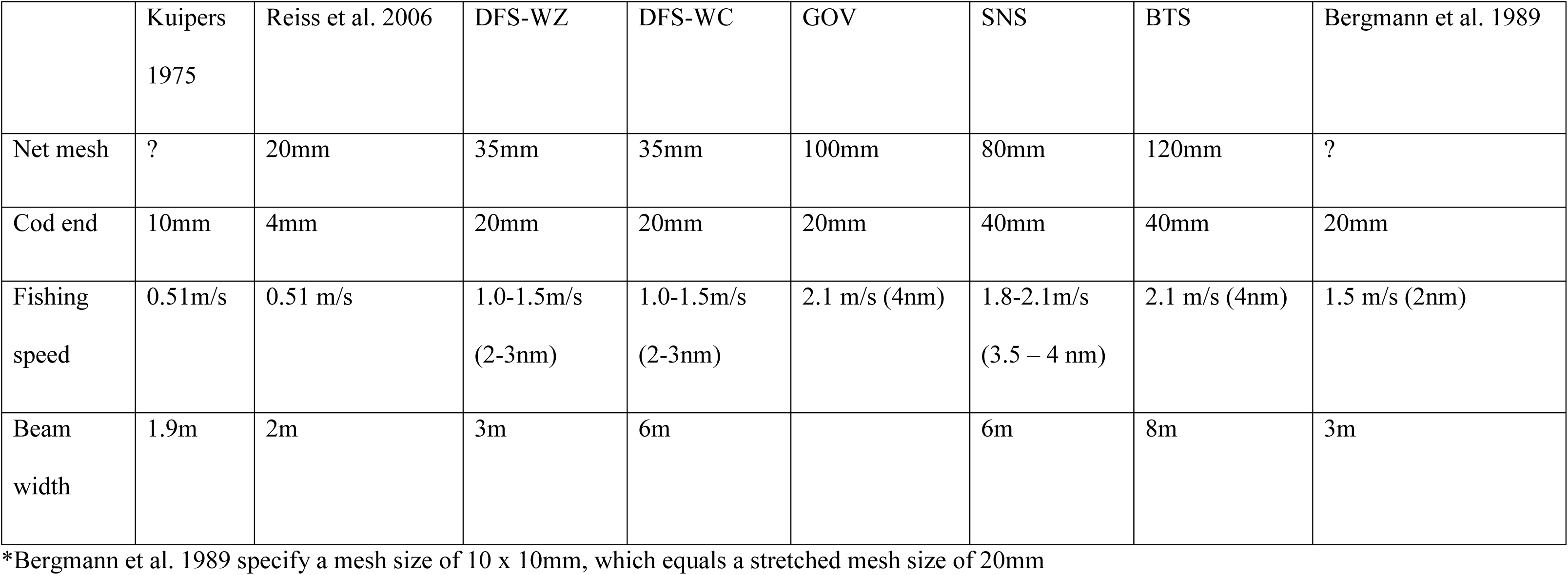
Characteristics of the different survey gear types -

|  | Kuipers<br>1975 | Reiss et al. 2006 | DFS-WZ | DFS-WC | GOV | SNS | BTS | Bergmann et al. 1989 |
| --- | --- | --- | --- | --- | --- | --- | --- | --- |
| Net mesh | ? | 20mm | 35mm | 35mm | 100mm | 80mm | 120mm | ? |
| Cod end | 10mm | 4mm | 20mm | 20mm | 20mm | 40mm | 40mm | 20mm |
| Fishing<br>speed | 0.51m/s | 0.51 m/s | 1.0-1.5m/s<br>(2-3nm) | 1.0-1.5m/s<br>(2-3nm) | 2.1 m/s (4nm) | 1.8-2.1m/s<br>(3.5 – 4 nm) | 2.1 m/s (4nm) | 1.5 m/s (2nm) |
| Beam<br>width | 1.9m | 2m | 3m | 6m |  | 6m | 8m | 3m |

\*Bergmann et al. 1989 specify a mesh size of 10 x 10mm, which equals a stretched mesh size of 20mm

**Table S2.** Caloric density of the 10 main prey items found in the harbour seal diet.

| Species/Family | Latin name | Caloric<br>density<br>(kcal/g) | Range | Source |
| --- | --- | --- | --- | --- |
| European flounder | *Platichthys flesus* | 1.09 | 0.99-1.20 | (Summers 1979)<sup>†</sup> |
| Sandeel | *Ammodytes sp.* | 1.41 | 0.88-1.48 | (Pedersen and Hislop 2001)<sup>*</sup> |
| Dover sole | *Solea solea* | 0.91 | 0.61-1.20 | (Soriguer et al. 1997)<sup>‡</sup> |
| Five-bearded rockling | *Ciliata mustela* | 1.02 | 0.88-1.29 | Average of round fish; cod and whiting |
| Whiting | *Merlangius merlangus* | 1.10 | 0.91-1.29 | (Pedersen and Hislop 2001)<sup>*</sup> |
| European Plaice | *Pleuronectes platessa* | 0.87 | 0.61-0.87 | (Costopoulos and Fonds 1989)<sup>§</sup> |
| Atlantic cod | *Gadus morhua* | 0.94 | 0.88-1.05 | (Härkönen and Heide- Jørgensen 1991) |
| Common dragonet | *Callionymus lyra* | 1.02 | 0.88-1.29 | Average of round fish; cod and whiting |
| Common dab | *Limanda limanda* | 0.87 | 0.61-0.87 | Assumed similar to Plaice |
| Bull-rout | *Myoxocephalus scorpius* | 1.02 | 0.88-1.29 | Average of round fish; cod and whiting |

<sup>†</sup>based on the average caloric density for 0 and I group flounder measured in July-Sept of 4.6 kcal/g dry weight. For July, the relationship between wet weight (x, in g) and dry weight (y, in g) was defined as $y = -0.36 + 0.24x$ (Summers 1979). Therefore, the wet-dry weight conversion factor for a 100 g flounder was 0.236, resulting in an estimated energy density of 1.09 kcal/g wet weight.
\*Converted from kJ/g to kcal/g using the conversion factor of 0.239. Caloric density estimates were based on fish >10cm caught in quarter 3.
Range was based on minimum and maximum energy density observed in all size-classes and seasons.
§In the Wadden Sea, the condition factor varied between 0.75 in winter, and 0.95 in summer, which corresponds to caloric densities between 0.61-0.87 (Costopoulos & Fonds 1989).
‡ Range based on minimum and maximum value of the range for plaice and flounder.

**Figure S1.**
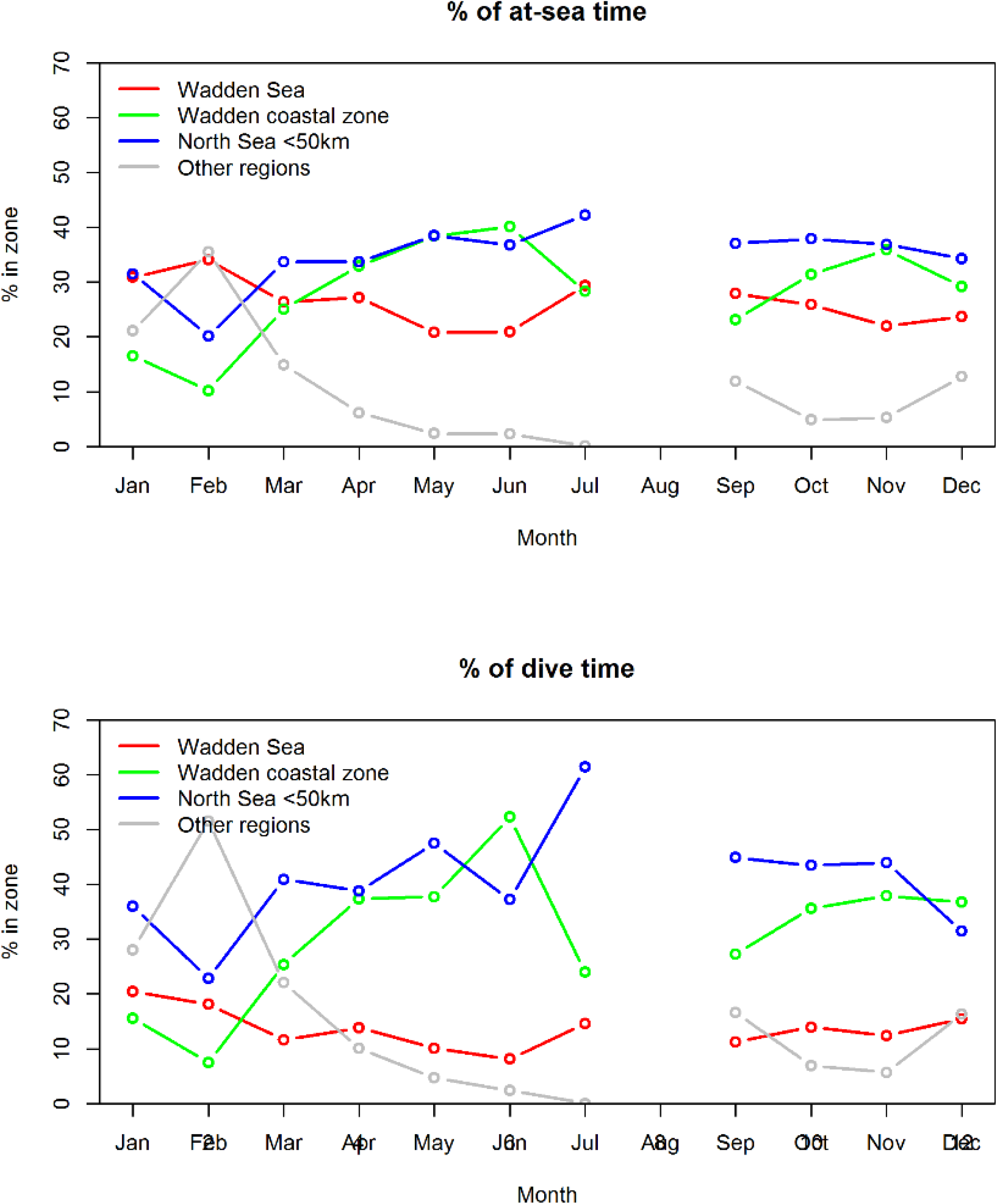
Proportion of time spent (top figure) and proportion of dive time (< −1.5m, bottom figure) in the different regions

**Figure S2.**
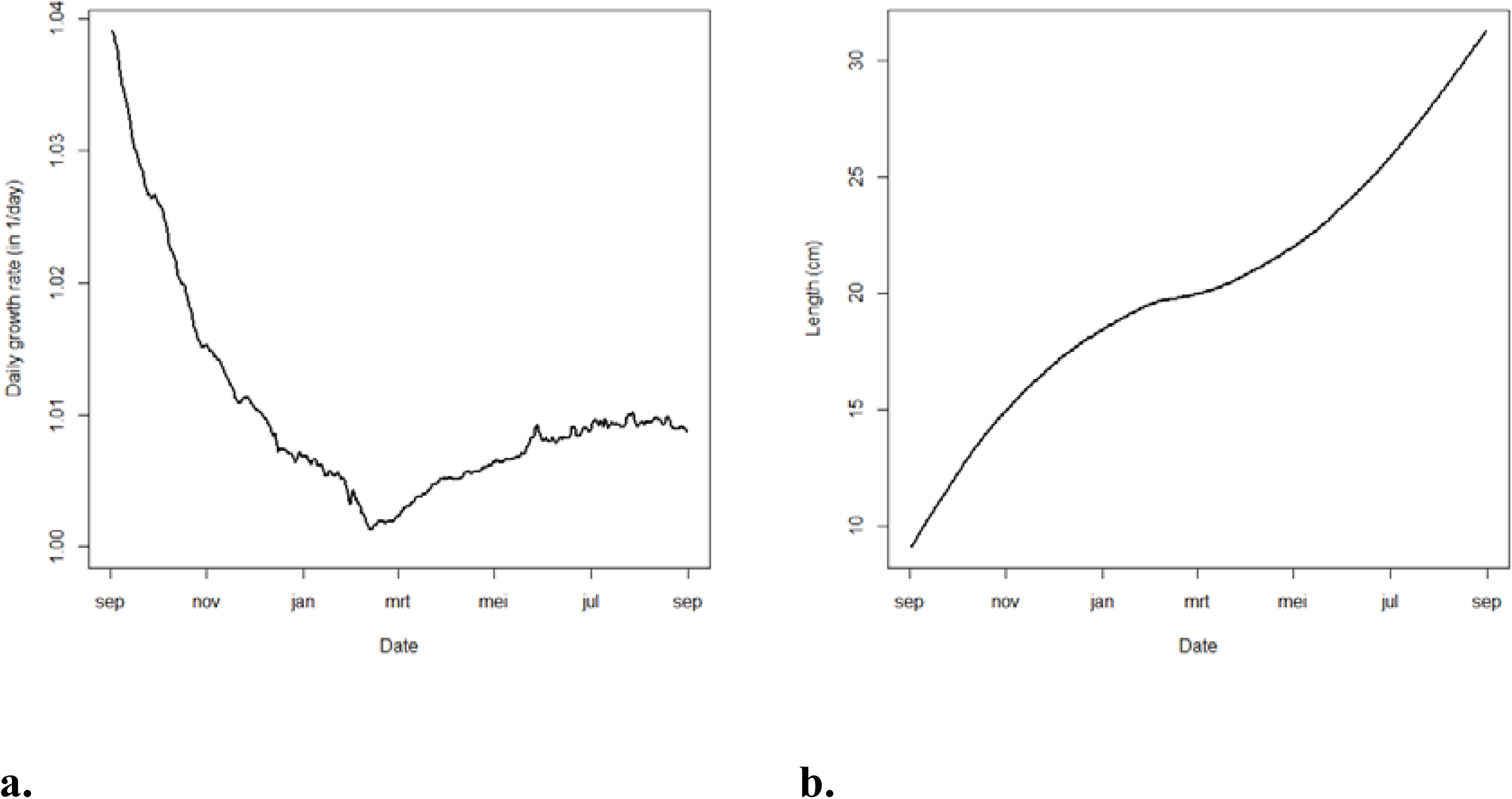
Growth of 0-group plaice per day based on *optimal* DEB model conditions (**a**), and how this leads to the change in length (in cm, **b**) and weight (in gram, **c**) throughout the year.

**Figure S3a-e.**
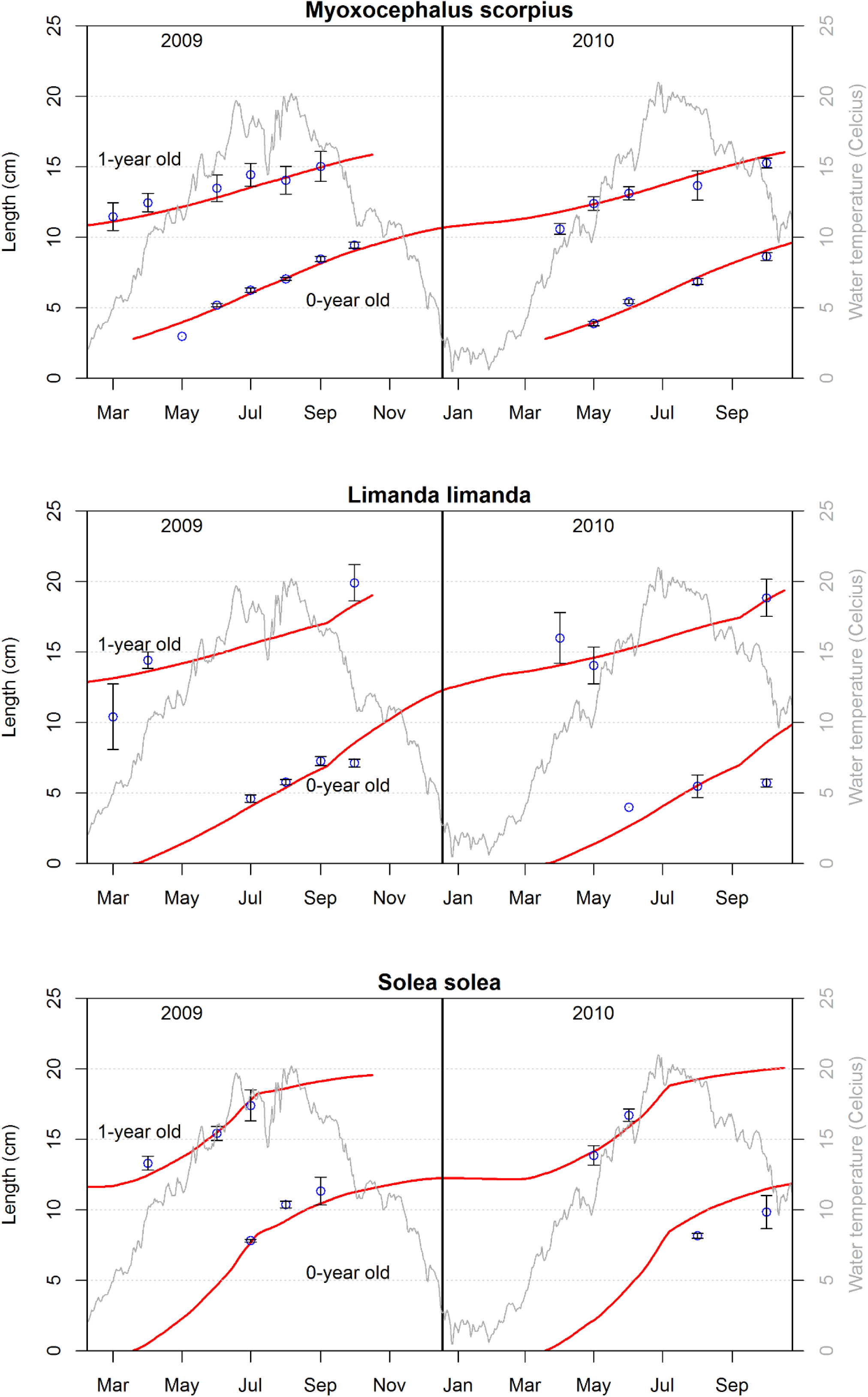
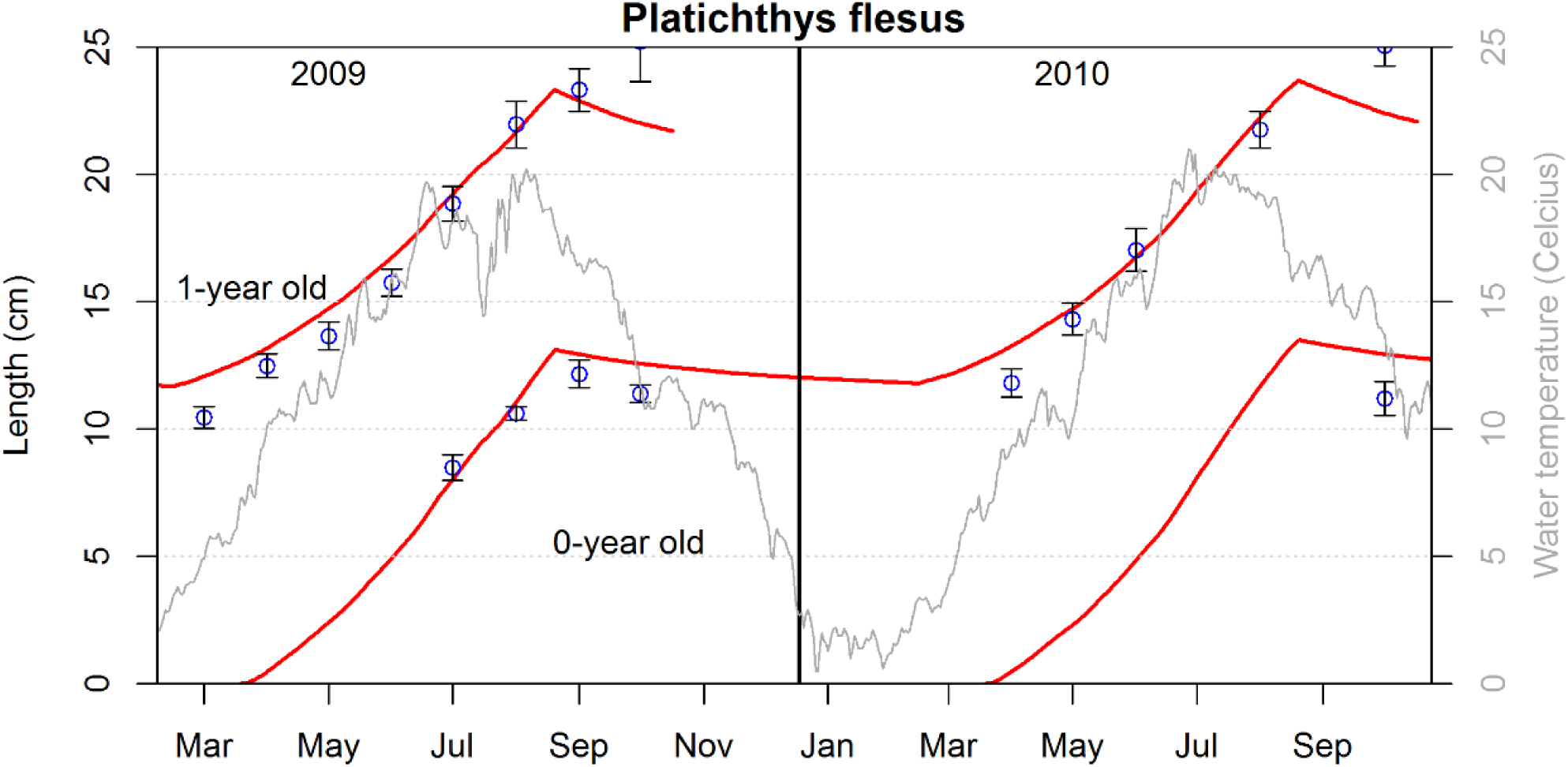
Observed length (circles and standard error bars) and predicted length (solid red line) of 0-and 1-year old flounder (*Platichthys flesus*), bull-rout (*Myoxocephalus scorpius*), dab (*Limanda limanda*) and sole (*Solea solea*). Grey line indicates water temperature. Predicted growth (solid lines) is based on DEB model where food intake parameter is allowed to vary between the winter months and other times of the years.

**Figure S4.**
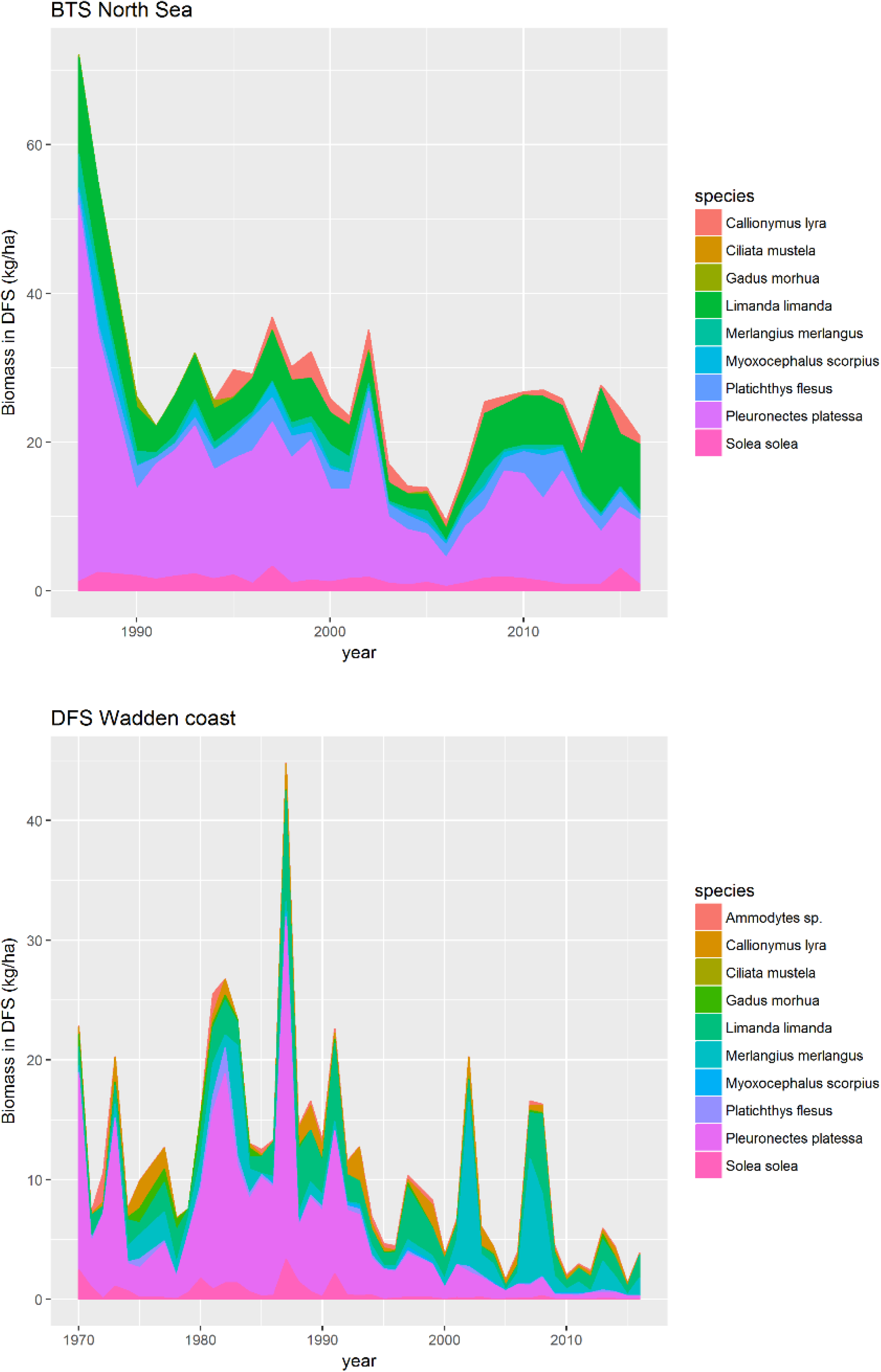
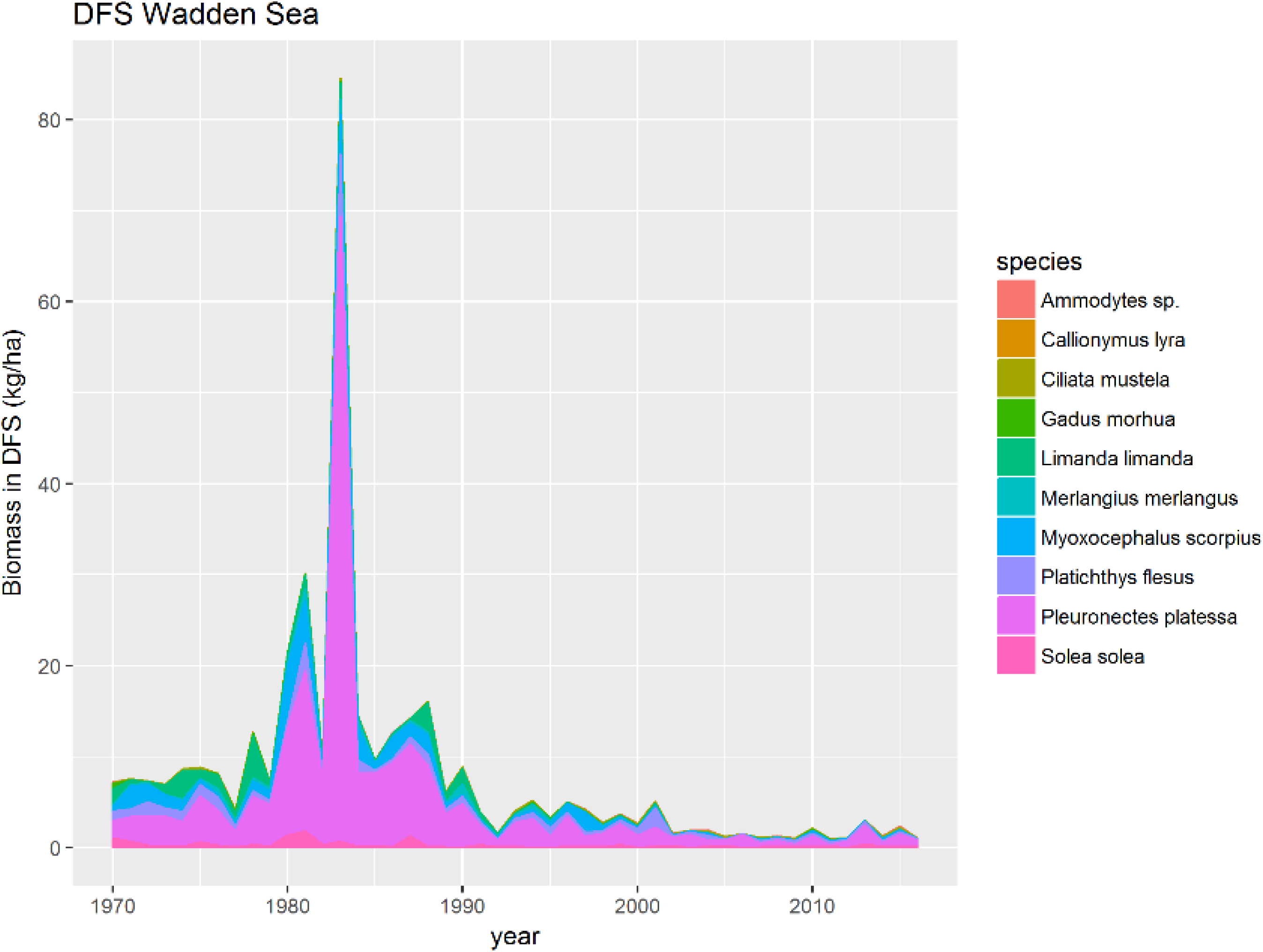
Species composition. Biomass (kg/ha) in BTS (top figure) and DFS Wadden coast and Wadden Sea (bottom figure)

**Figure S5.**
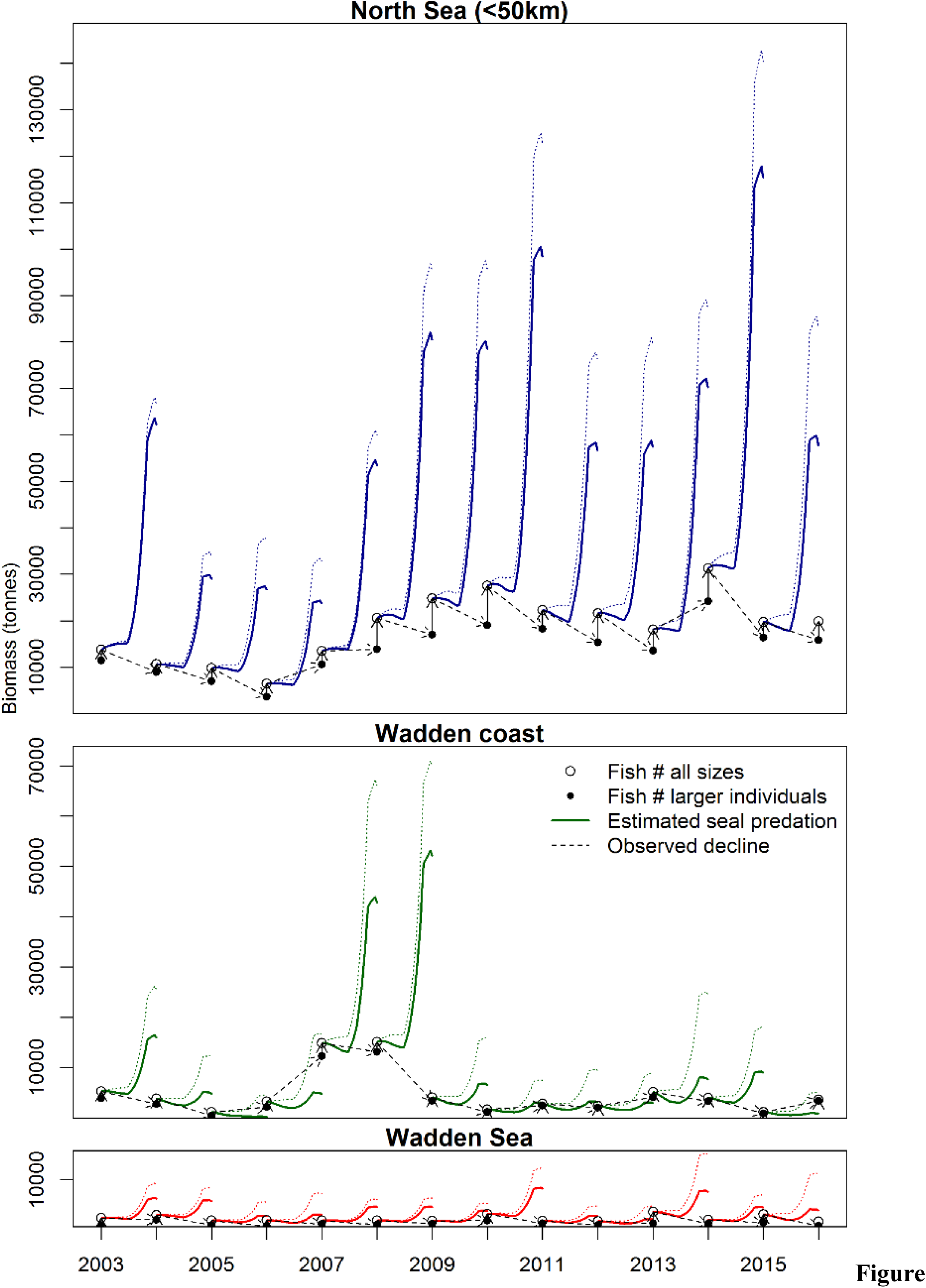
The estimated effect of seal predation on the biomass of prey fish for the Wadden Sea (red lines), Wadden coast (green lines) and remaining areas up to 50km from the haul-outs (blue lines, and see regions in Figure 1). The fish survey in September (i.e. BTS for the North Sea offshore zone up to 50km from the nearest haul-out, and DFS for the Wadden coast and Wadden sea) is the starting point (open circle), after which biomass decline due to seal predation. During the summer season, a rapid increase in biomass occurs, except in years when the number of individuals has drastically reduced. In the subsequent survey, the biomass of remaining fish (i.e. larger individuals, >13.5cm) has declined, but is again supplemented with new recruits.

